# DNA and H3K4 methylation act cooperatively to protect regulatory elements

**DOI:** 10.64898/2026.11.12.766180

**Authors:** Angélique David, Marlet Morales-Franco, Narjesse Hounani, Ana Monteagudo-Sánchez, Suzy Perdrix, Julien Richard Albert, Margherita Scarpa, Beyza Bozdag, Thomas Damagnez, Maxim V.C. Greenberg

## Abstract

Histone H3 lysine 4 methylation (H3K4me) is associated with transcriptionally active regulatory elements: H3K4me1 is typically found at enhancers, and H3K4me2/3 at gene promoters. In mammals, the presence of H3K4me can reinforce expression programs, but it might also help protect from epigenetic silencing complexes. To wit, H3K4me antagonizes the binding of the *de novo* DNA methyltransferase enzymes, which catalyze the repressive 5-methylcytosine (5meC) epigenetic mark. To determine the extent to which H3K4me protects regulatory elements from 5meC deposition—and how this influences genome regulation—we used mouse embryonic stem cells (ESCs), which are resilient to the complete absence of 5meC. Employing an inducible dominant negative transgene that globally reduces H3K4me, we assessed the impact in both presence and absence of 5meC. Surprisingly, while ESCs could withstand reduced H3K4me or 5meC, their combined loss led to catastrophic cellular phenotypes in pluripotency and during differentiation. Genomic analyses revealed that H3K4me and 5meC work in concert to protect regulatory elements from Polycomb-mediated silencing. Thus, this work unraveled an unexpected functional cooperation between two epigenetic marks typically viewed as antagonistic in safeguarding transcriptional programs.

## Introduction

In mammals, active regulatory elements are characteristically enriched for histone H3 lysine 4 methylation (H3K4me). H3K4 monomethylation (H3K4me1) is frequently found at enhancers, while H3K4 di- and trimethylation (H3K4me2/3) are deeply conserved promoter marks (Wang and Helin 2024). The function of H3K4me at both enhancer and promoter regions remains an active area of research (Howe et al. 2017). H3K4 methyltransferases are large enzymes assembled in multiprotein complexes and exert effects that are not directly related to catalytic activity (Morgan and Shilatifard 2023). At promoters, some studies have demonstrated that H3K4me3 can be instructive for gene activation (Anyetei-Anum et al. 2024; Lauberth et al. 2013; Policarpi et al. 2024), while others suggest it is unnecessary (Clouaire et al. 2014; Hughes et al. 2023; Au et al. 2025). This paradox also might reflect context-specific roles of H3K4me3 depending on the loci examined. The function of H3K4me1 is even more confounding, as very few chromatin reader proteins associated with activation have been demonstrated to bind to the mark (Lukauskas et al. 2024).

A potential function of H3K4me that is not mutually exclusive with a role in activation, *per se*, is protection from 5-cytosine DNA methylation (5meC)-mediated silencing. 5meC predominantly occurs in the CpG dinucleotide context and is therefore referred to interchangeably with CpG methylation throughout this text. The *de novo* DNA methyltransferase 3A and 3B (DNMT3A/B) enzymes contain an ADD domain that binds to unmethylated H3K4 residues, allowing the catalytic domain to access and methylate DNA. H3K4me, especially H3K4me2/3, inhibits this binding, rendering DNMT3A/B inactive (Ooi et al. 2007; Guo et al. 2015). Most promoters in mammalian genomes, including almost all housekeeping and developmental genes, exhibit a so-called “CpG Island” (CGI) profile, which is characterized by high CpG density and near absence of 5meC (Greenberg and Bourc’his 2019). While H3K4me2/3 is not the sole reason that CGI promoters are faithfully protected from 5meC deposition, it could certainly play an important role (Hughes et al. 2020). Consistently, during early mouse development, failure to mark certain promoters with H3K4 methylation leads to 5meC-mediated silencing that results in lethality (Eckersley-Maslin et al. 2020; Gretarsson and Hackett 2020). The relationship between DNA and H3K4 methylation is less clear at enhancers. Although 5meC is generally anticorrelated with enhancer activity (Charlton et al. 2020; Stadler et al. 2011), recent work from our lab suggests that H3K4me1 and 5meC are not mutually exclusive marks, and 5meC levels are high (≥50% methylated CpGs) at a substantial proportion of active enhancer elements (Morales-Franco et al. 2026).

In order to interrogate the relationship between H3K4me and 5meC, a developmentally relevant model is mouse embryonic stem cells (ESCs), which are pluripotent and can differentiate into all somatic cell types. Crucially, ESCs can tolerate complete absence of 5meC in the triple knockout (TKO) of *Dnmt3a/b* and the maintenance methyltransferase, *Dnmt1* (Tsumura et al. 2006). While there are six major H3K4 methyltransferase complexes, the degradation of shared core complex components leads to absence of all forms of H3K4me (Wang et al. 2023; Hu et al. 2023). However, this results in severe growth and self-renewal defects in ESCs. An alternative system is an inducible version of the replication-independent histone variant H3.3 harboring a lysine-to-methionine (K-to-M) mutation at position 4 (H3K4M). H3K4M acts as a dominant negative, leading to global reduction of H3K4me, but is better-tolerated by ESCs (Chan et al. 2013; Lewis et al. 2013; Jang et al. 2017, 2019; Warren et al. 2024; Yagi et al. 2025).

We introduced the H3K4M transgene into both wild-type (WT) and TKO ESCs, with the hypothesis that 5meC-mediated silencing observed in the WT H3K4M line following H3K4me depletion would be suppressed in the TKO H3K4M cells. Surprisingly, we instead observed that while WT H3K4M ESCs appeared healthy, TKO H3K4M cells exhibited a severe proliferation phenotype and displayed drastic differentiation defects correlated with pronounced transcriptional dysregulation. We went on to demonstrate that both H3K4me and 5meC cooperatively protect regulatory elements from H3K27me3 deposition by Polycomb Repressive Complex 2 (PRC2). Unrestrained PRC2 activity in TKO H3K4M led to increased silencing at enhancers and promoters, likely explaining the cellular phenotypes. Finally, we performed epigenome editing to probe the functional interplay between these three axes of epigenetic control at selected promoters, providing further support for our interpretations of the genome-wide data. In sum, our findings show a novel cooperative role that H3K4me and 5meC play in genome regulation and cell-fate transition—two marks that are traditionally considered as antagonistic.

## Results

### H3K4M induction leads to H3K4me depletion and cellular defects in pluripotent cells

To investigate the interplay between H3K4me and 5meC, we inserted a doxycycline (Dox)-inducible H3K4M transgene at the *Tigre* safe harbor locus in mouse ESCs (Zeng et al. 2008). As a control, we established independent lines carrying a Dox-inducible unmodified histone H3.3 transgene (hereafter: H3) targeted to the same locus. The transgenes were introduced into *Dnmt* WT and TKO backgrounds (Fig. 1A). Mass spectrometry analysis revealed an increased fraction of unmodified H3K4 peptides (H3K4me0) in H3K4M-expressing lines relative to their respective H3 controls after 24 hours (h) of Dox induction (Fig. S1A). Moreover, a substantial reduction in H3K4me1 and H3K4me2 levels in both H3K4M lines was observed. In contrast, the decrease in H3K4me3 levels was more variable and did not reach statistical significance, although it should be noted that this is consistent with data generated from a study using a similar approach (Yagi et al. 2025) (Fig. S1A). In addition, in line with previous reports (King et al. 2016), TKO H3 cells displayed higher levels of H3K4me than WT H3 cells, although the increase reached statistical significance only for H3K4me1. Depletion of H3K4me in H3K4M lines was further confirmed by immunoblot analysis after 48 h of transgene expression, showing that all three H3K4me states were similarly reduced in WT and TKO H3K4M-expressing cells, but not in the corresponding H3 control lines, nor in the uninduced cell lines (Fig. 1B). Notably, H3K4me3 levels were markedly decreased in both H3K4M-induced lines, confirming the reduction of this mark despite ambiguous mass spectrometry results. Importantly, transgene expression levels were comparable across cell lines. Therefore, we established an inducible H3K4M-based system that enables efficient depletion of all H3K4 methylation marks within 48 h of induction. This experimental framework allows for the analysis of H3K4me and 5meC dynamics in pluripotent stem cells that can be readily differentiated.

**Figure 1.**
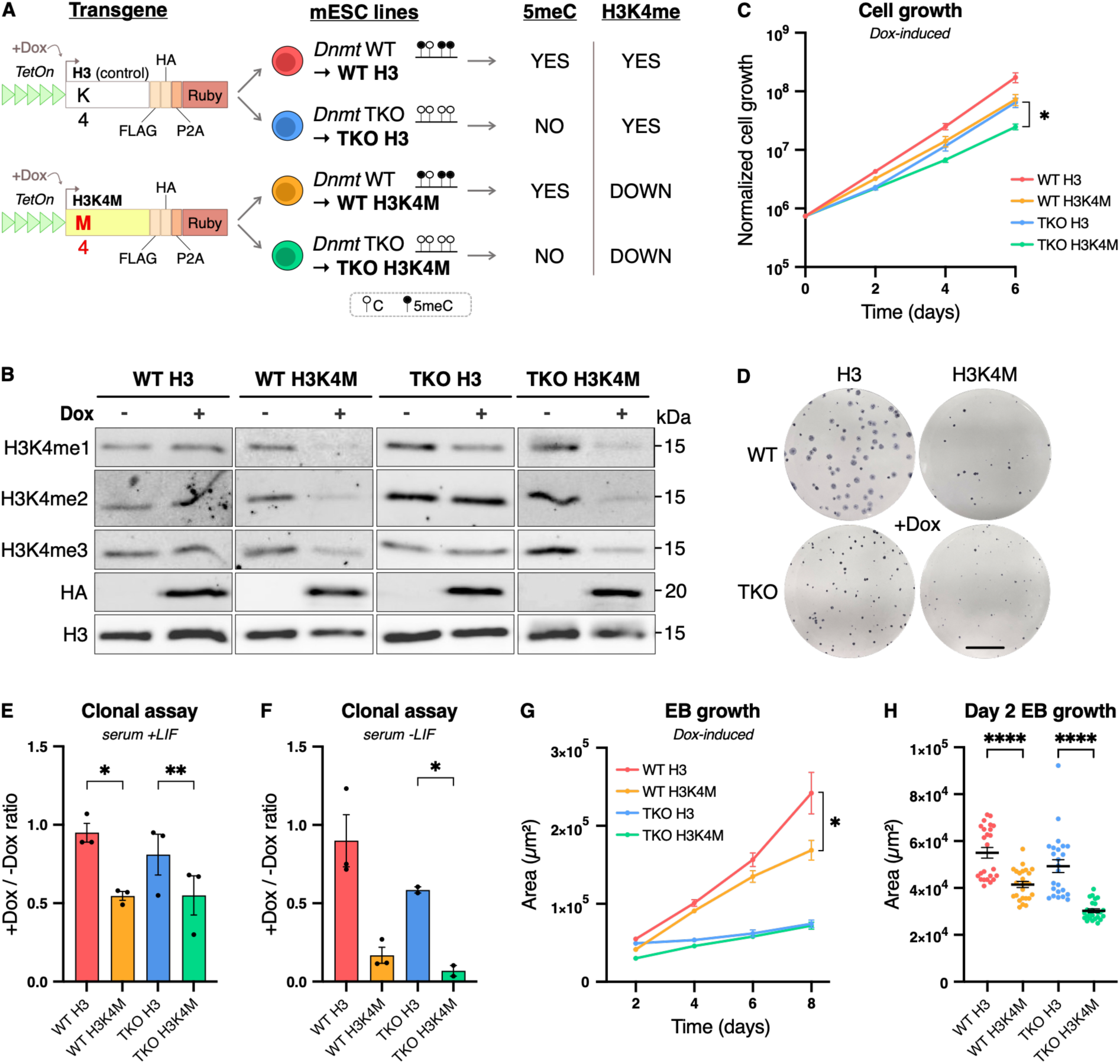
H3K4M transgene expression leads to H3K4me depletion and cellular defects in pluripotent and early-differentiated cells. **A.** Schematic of the inducible system used in this study. A wild-type H3.3 (H3) histone transgene serves as a control (top). The H3K4M construct harbors a lysine-to-methionine (K-to-M) substitution at the fourth amino acid residue (bottom). Transgene expression is driven by a TetOn promoter and induced upon doxycycline (Dox) treatment, resulting in global H3K4me depletion. Both constructs include FLAG and HA tags and a P2A-linked Ruby reporter to monitor transgene expression. Transgenes were inserted at the Tigre safe-harbor locus, in *Dnmt* WT or TKO cells, the latter being devoid of 5meC. **B.** Representative immunoblot showing H3K4me (H3K4me1/2/3) levels and transgene expression (HA) in uninduced (-Dox) or induced (+Dox, 48 h) cell lines grown in serum+LIF. C. Six-day growth assay in serum+LIF +Dox culture conditions. Normalized cell growth is represented in Iog10 scale, n = 3; except for WT H3, n = 5. D. Representative images of alkaline phosphatase-stained colony formation assays in serum+LIF +Dox conditions. The scale bar corresponds to 1 cm. E, F. Ratio of alkaline phosphatase-stained colony numbers in induced (+Dox) over uninduced (-Dox) serum+LIF (E) or serum-LIF (F) conditions for all cell lines. Replicates and their mean are shown, n = 3. **G, H.** Embryoid body (EB; +Dox) growth as measured by area (pm2) over 8 days of differentiation (G) and at day 2 of differentiation (H). n ≥ 24 at each time point, obtained from two independent experiments. **C, E, F, G, H:** Data are shown as mean ± s.e.m. of replicates from independent experiments. P-values were calculated using a paired Student’s t-test **(E, F)** or an unpaired Student’s t-test **(H),** on day 6 normalized cell counts **(C)** and on day 8 measurements (G). * P≤0.05, ** P≤0.01, **** P≤0.0001.

Previous reports have shown that H3K4me depletion following H3K4M expression leads to non-significant proliferation defects in WT ESCs (Jang et al. 2017). Moreover, TKO cells exhibit a proliferation rate comparable to that of WT cells (Tsumura et al. 2006). Thus, H3K4me and 5meC are individually largely dispensable for ESC growth. To investigate the effects of H3K4me loss in a 5meC-absent background on pluripotent cell proliferation, a six-day growth assay was performed on independent clonal lines in fetal bovine serum (serum) + Leukemia Inhibitory Factor (LIF) conditions (Fig. 1C; Fig. S1B) (Graf et al. 2011). Notably, when cultured in serum+LIF, WT cells harbor elevated 5meC levels, with ∼70-80% of methylated CpGs (Habibi et al. 2013; Ficz et al. 2013; Azevedo Portilho et al. 2021), allowing for the comparison of cellular phenotypes between high (WT) and absent (TKO) 5meC contexts. In non-induced conditions, cells showed very little difference in growth (Fig. S1B). Upon Dox induction, both WT H3K4M and TKO H3 cells displayed a mild but non-significant decrease in proliferation compared to WT H3, consistent with published results (Jang et al. 2017; Tsumura et al. 2006) (Fig. 1C). Strikingly, TKO H3K4M cells exhibited more pronounced growth defects compared to the TKO H3 control (Fig. 1C). Rather than rescuing the mild proliferation defects observed in WT H3K4M or TKO H3, loss of H3K4me in the TKO background further exacerbated the defective growth phenotype, with an additive effect compared to perturbation of either modification alone (Fig. S1C).

Next, to test whether the depletion of one or both epigenetic modifications affects ESC self-renewal and maintenance of the pluripotent state, we performed an alkaline phosphatase-stained colony formation assay. Previous studies have shown that 5meC is dispensable for ESC self-renewal (Tsumura et al. 2006; Shukla et al. 2020). Uninduced WT lines, as well as Dox-induced WT H3 controls, displayed heterogeneous colony morphology (Fig. 1D; Fig. S1D,E). A majority (∼80%) of colonies displayed partial alkaline phosphatase staining (hereafter referred to as “mixed”), whereas a minority (∼20%) consisted of smaller, rounder, and more intensely stained colonies associated with undifferentiated cells. TKO H3-induced and uninduced TKO colonies resembled the latter category, likely indicating their incapacity for differentiation (Sakaue et al. 2010). Importantly, both WT and TKO H3K4M-induced lines exhibited significant colony-forming defects, producing fewer and smaller colonies (Fig. 1D; Fig. S1E), with a significant reduction in the induced-to-uninduced colony number ratio compared to control lines (Fig. 1E). Notably, H3K4me loss appeared more detrimental to WT cells at low density than in a population-growth assay. Together, these observations indicate that H3K4me depletion similarly alters self-renewal and colony-forming capacity in both WT and TKO ESCs.

To independently validate that the defects observed in H3K4M lines are due to H3K4me depletion, we generated WT and TKO RbBP5-FKBP degron lines, as previously described for WT ESCs (Wang et al. 2023; Au et al. 2025) (Fig. S1F). As part of the “WRAD” set of proteins present in all the major H3K4 methyltransferase complexes, RbBP5 is essential for H3K4me deposition (Zhang et al. 2015), and its degradation upon dTAG-13 addition leads to nearly complete elimination of H3K4me (Wang et al. 2023). RbBP5-depleted cells exhibited a significant decrease in proliferation compared to control lines in a serum growth assay (Wang et al. 2023) (Fig. S1G). Importantly, RbBP5 degradation in TKO cells resulted in more severe proliferation defects than in WT cells by day 6 (Fig. S1G,H). Additionally, a colony formation assay with alkaline phosphatase staining showed a 2.5-fold reduction in colony number for WT RbBP5-depleted cells compared to control cells (Fig. S1I,J). TKO RbBP5-depleted cells exhibited an even greater impairment in self-renewal, with almost no colony formation after dTAG-13 addition (Fig. S1I,J). Overall, the proliferation and self-renewal defects observed in H3K4M cells were consistent but more severe in the RbBP5-FKBP degron system following RbBP5 degradation. In particular, RbBP5 depletion in absence of 5meC resulted in exacerbated defects compared to a WT context.

Finally, to distinguish genetic from epigenetic effects in the TKO H3K4M line following H3K4me loss, we used a DNMT1 inhibitor (GSK-3484862; DNMT1i (Azevedo Portilho et al. 2021)) to acutely deplete 5meC in WT H3 and H3K4M cells. Reduction of 5meC was confirmed using a LUminometric Methylation Assay (LUMA) (Fig. S1K): WT cells grown for 4 days in the presence of DNMT1i retained ∼20% 5meC, compared to an average of ∼70% in control cells. Despite incomplete 5meC removal, markedly aggravated proliferation defects were observed in WT H3K4M +DNMT1i compared to WT H3 +DNMT1i cells (Fig. S1L). Importantly, the dual depletion of 5meC and H3K4me marks led to a synergistic effect on cell growth (Fig. S1M). These results support the conclusion that defects observed in TKO H3K4M line are mainly caused by the combined absence of 5meC and the acute loss of H3K4me. Together, these data demonstrate that H3K4me is required for robust proliferation and self-renewal in serum-grown ESCs, and the effect worsens in the absence or reduction of DNA methylation. This reveals a surprising functional interplay between these two epigenetic modifications, given that they are widely presumed to be functionally antagonistic.

### H3K4M-associated defects are exacerbated during early differentiation

Having established that H3K4me is necessary for robust ESC self-renewal, and, together with 5meC, contributes to proper cell proliferation, we next wondered how the loss of one or both marks would affect cells’ capacity to differentiate. First, to test single cells’ ability to survive and expand while exiting pluripotency, a colony formation assay followed by alkaline phosphatase staining was performed in the absence of LIF, removal of which promotes spontaneous differentiation (Cherepkova et al. 2016). Both H3K4M lines showed colony-forming defects in -LIF that were exacerbated in the TKO H3K4M line (Fig. 1F; Fig. S2A). These results were again confirmed with the RbBP5-FKBP degron lines: RbBP5-depleted WT and TKO lines displayed a dramatic decrease in colony numbers compared to untreated cells, with almost no colonies surviving (Fig. S2B), consistent with the more severe phenotypes observed in RbBP5-depleted lines compared to H3K4M lines. These results demonstrate that undirected differentiation is strongly impaired in H3K4me-depleted cells, regardless of 5meC presence.

Next, embryoid body (EB) differentiation was conducted to more precisely assess the effects of loss of H3K4me and 5meC, separately or together, on early differentiation. Original work reported that TKO cells display growth delays and increased apoptosis during EB formation: they can form three-dimensional structures but not proper EBs, due to the requirement for 5meC to establish stable consolidation of the developmental program during differentiation (Tsumura et al. 2006; Sakaue et al. 2010; Schmidt et al. 2012; Schulz et al. 2024). In our cell line system, WT H3K4M EBs formed properly but exhibited a smaller size compared to WT H3 EBs, from day 2 of differentiation, with the difference becoming even more pronounced by day 8 (Fig. 1G; Fig. S2C). TKO H3 and H3K4M EBs could form spheroids, but barely grew past day 2, as expected (Schulz et al. 2024). Importantly, at day 2 of differentiation, we observed a significant reduction in EB size in both H3K4M lines compared to respective control lines, with TKO H3K4M EBs being the most affected (Fig. 1H).

By phenotypic assessment, H3K4me appears to play a role during the earliest stages of differentiation without being strictly required for EB formation, whereas 5meC depletion masks H3K4me-associated phenotypes at later stages, consistent with its known role in sustaining developmental programs. Nevertheless, 5meC and H3K4me are concomitantly required during early differentiation to sustain EB growth.

### Absence of 5meC exacerbates H3K4M-associated transcriptional defects

Next, to investigate the molecular bases underlying these phenotypes, we assessed how H3K4M expression impacts transcription during EB differentiation in H3K4M lines. By RT-qPCR, we observed transgene silencing in WT cells from day 2 onward, whereas expression in TKO cells increased and stabilized by day 4 (Fig. S2D). Given the discrepancies in transgene expression, we performed RNA-seq in ESCs (day 0) and EBs (day 2), a timepoint at which transgene RNA levels across cell lines were the most homogeneous (Fig. S2D,E), and at which reduced EB size was also observed in both H3K4M lines (Fig. 1H). Principal component analysis (PCA) showed that samples clustered primarily by differentiation stage and genetic background (WT versus TKO); although the H3K4M transgene did exert an effect, particularly at day 2 (Fig. 2A). To further assess the impact of H3K4me depletion on transcription, we performed pairwise comparisons to identify differential gene expression between H3 and H3K4M lines within the same genetic background. No differentially expressed genes (DEGs) were detected on day 0 WT H3K4M ESCs (Fig. S3A), consistent with the minimal effect observed in the serum growth assay. In contrast, TKO H3K4M ESCs displayed 178 DEGs relative to TKO H3, with a majority (73%) of them downregulated (Fig. S3B). Gene Set Enrichment Analysis (GSEA) (Mootha et al. 2003; Subramanian et al. 2005) revealed distinct pathways depending on the genetic background, with immune-related terms specific to TKO H3K4M, and a set of developmental processes specific to the WT H3K4M cells, or shared between both H3K4M lines (Fig. S3C). Notably, most genes downregulated in TKO H3K4M showed higher expression in TKO H3 relative to all cell lines (Fig. 2B), suggesting that loss of H3K4me in a 5meC-depleted context counteracts gene upregulation observed upon removal of 5meC-dependent repression. Curiously, a similar trend was observed for genes upregulated in TKO H3K4M, which tended to be downregulated in TKO H3 (Fig. S3D). In addition, for the same set of TKO H3K4M DEGs, WT H3K4M cells showed similar but weaker expression changes compared to WT H3 control, indicating modest transcriptional effects of H3K4me depletion in this context (Fig. 2B; Fig. S3D). Overall, short-term H3K4me loss in WT pluripotent cells does not lead to major transcriptomic changes. In contrast, in TKO H3K4M cells, the additional 5meC absence is correlated with stronger transcriptional alterations, predominantly gene downregulation.

**Figure 2.**
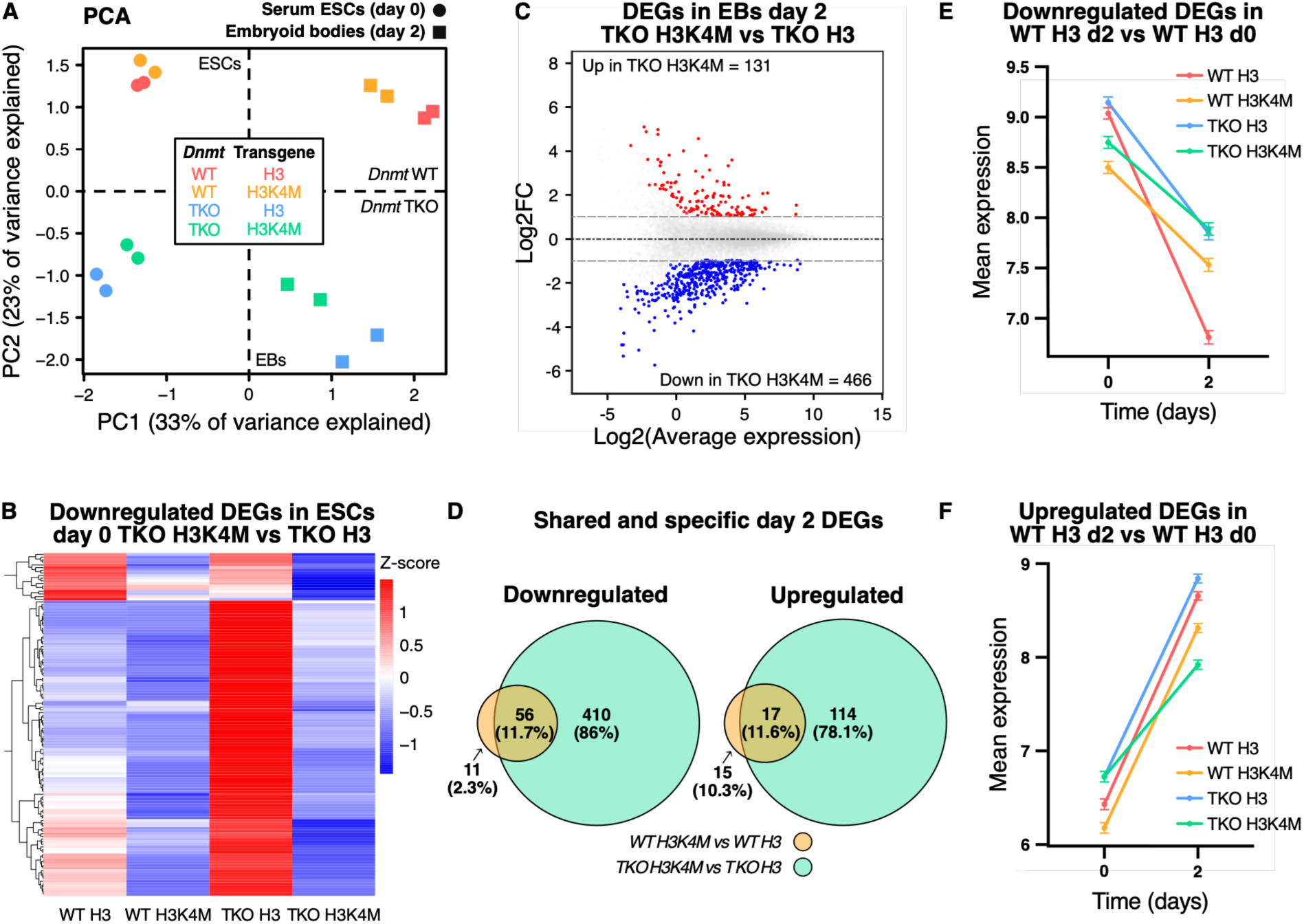
Absence of 5meC aggravates H3K4M-induced transcriptomic defects. **A.** Principal Component Analysis (PCA) of transcriptomic datasets representing ESCs grown in serum+LIF (day 0) and embryoid bodies (EBs; day 2) (+Dox). Each cell line at each time point is represented by two replicates. B. Heatmap of row-normalized gene expression (Z-score) for downregulated differentially expressed genes (DEGs (log2FC < -1 and adjusted p-value (t-test) < 0.05); n = 130) in the TKO H3K4M vs TKO H3 comparison in day 0 samples. The mean of two replicates is shown. **C.** MA plot of differential gene expression (|log2FC| > 1 and adjusted p-value (t-test) < 0.05) in the TKO H3K4M vs TKO H3 comparison in day 2 embryoid bodies. The average expression is represented as Iog2. Numbers of significantly up- and downregulated genes are indicated. D. Venn diagram representing the shared and specific number of down-(left) and upregulated (right) DEGs between WT H3K4M vs WT H3 and TKO H3K4M vs TKO H3 comparisons in day 2 embryoid bodies. **E,** F. Mean expression levels (log2(RPKM)) for downregulated (E; n = 990) or upregulated (F; n = 1242) genes in WT H3 during differentiation (day 2 vs day 0 comparison) in all cell lines. Data are shown as mean ± sd of two replicates.

At day 2 of embryoid body differentiation, 99 genes were dysregulated in WT H3K4M compared to WT H3, among which 67.7% were downregulated (Fig. S3E). In the TKO background, we observed a sixfold increase in DEGs compared to the WT comparison, with 78% of genes being downregulated (Fig. 2C). Similarly to day 0, developmental pathways were enriched in WT H3K4M, whereas TKO H3K4M cells rather displayed enrichment of immunity-related pathways (Fig. S3F). Importantly, a major subset of WT H3K4M DEGs was shared with TKO H3K4M (downregulated = 56; upregulated = 17, Fig. 2D), suggesting that these genes are sensitive to H3K4me loss, while DEGs specific to TKO H3K4M likely reflect transcriptional changes resulting from the combined loss of 5meC and H3K4me.

Finally, we examined dynamic gene expression changes during differentiation by comparing DEGs obtained from day 2 vs day 0 pairwise comparisons within the same cell line. H3K4M lines, particularly TKO H3K4M cells, displayed fewer up- and downregulated DEGs across differentiation, possibly indicating impaired cellular transitions (Fig. S3G). To distinguish between defects in pluripotency exit and differentiation onset, we looked at the average expression levels of the up- or downregulated genes that occur in WT H3 during early differentiation from day 0 to day 2. First, genes downregulated in WT H3 were less efficiently repressed in other lines, particularly in the TKO background (Fig. 2E), indicating impaired exit from pluripotency in both 5meC- and/or H3K4me-depleted contexts. Conversely, genes upregulated in WT H3 during differentiation showed similar induction across all lines except in TKO H3K4M cells (Fig. 2F), indicating an apparently stronger defect in activating early differentiation programs in this context. This was further supported by the expression patterns of pluripotency markers, which were mainly expressed at lower levels in H3K4M lines at day 0 and not properly repressed at day 2—an effect aggravated in the TKO H3K4M line (Fig. S3H). On the contrary, in both H3K4M lines, early-differentiation genes showed reduced upregulation at day 2, particularly in TKO H3K4M EBs (Fig. S3I).

In conclusion, both H3K4me and 5meC contribute to proper exit from pluripotency, while the establishment of differentiation-specific transcriptional programs relies on the concerted interplay of both marks. However, an explanation for the apparent cooperation between H3K4me and 5meC remains lacking.

### PRC2 inhibition suppresses TKO H3K4M proliferation defects and partially rescues gene downregulation

The pervasive gene downregulation observed in the TKO H3K4M compared to TKO H3 challenged our expectations, given that H3K4me depletion in WT H3K4M line led to minor transcriptional changes, while 5meC absence in the TKO background is associated with global gene derepression. We therefore hypothesized that the TKO H3K4M line might be particularly susceptible to an additional layer of repression. A strong candidate is Polycomb-mediated silencing; multiple Polycomb group proteins are 5meC-sensitive (Farcas et al. 2012; Blackledge et al. 2014; Li et al. 2017; Perino et al. 2018), and Polycomb Repressive Complex 2 (PRC2) is antagonized by H3K4me (Schmitges et al. 2011; Hagarman et al. 2013; Wang and Helin 2024). To test this in serum-grown ESCs, we used a potent inhibitor of PRC2 activity (PRC2i) that targets the catalytic subunit, EZH2, and leads to a strong reduction in H3K27me3 levels (Fig. S4A).

Strikingly, PRC2 inhibition rescued the proliferation defects observed in TKO H3K4M cells (Fig. 3A). Notably, PRC2 inhibition also improved proliferation in TKO H3 cells; however, this effect was markedly more pronounced in the TKO H3K4M line, consistent with an enhanced sensitivity upon additional loss of H3K4me (Fig. 3B). This compelling result prompted us to investigate the impact of PRC2 inhibition on gene expression by RNA-seq. Addition of PRC2i led to a moderate increase of expression of the 130 genes that were downregulated in the TKO H3K4M line relative to TKO H3 (Fig. 3C,D). Importantly, this gene set was not globally impacted by PRC2i addition in the other cell lines. Examination of absolute expression levels (RPKM) revealed a slight decrease in TKO H3 upon PRC2i treatment, while expression increased in TKO H3K4M (Fig. S4B), supporting the context-dependent effect of PRC2 inhibition. In contrast, a set of randomly selected genes exhibited uniform effects across +PRC2i and -PRC2i conditions (Fig. S4C). Overall, these results suggest that the depletion of both 5meC and H3K4me creates a chromatin environment in which a subset of genes is vulnerable to PRC2-mediated repression. These dynamics may help explain the aggravated cellular phenotypes observed in the TKO H3K4M line.

**Figure 3.**
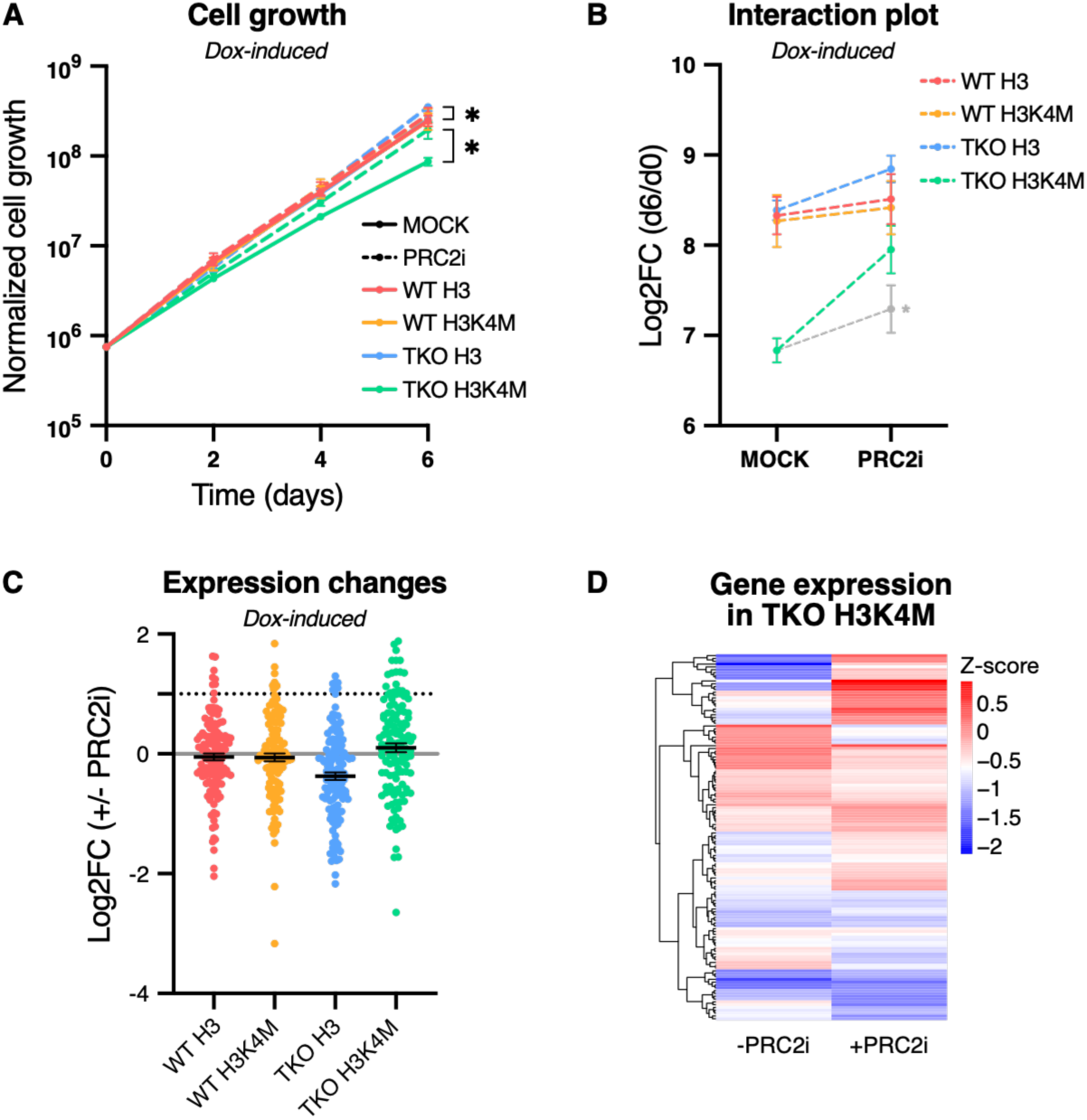
PRC2 inhibition suppresses TKO H3K4M proliferation phenotype in ESCs. **A.** Six-day growth assay in serum+LIF +Dox culture conditions. Cell lines are grown either in presence of a Mock inhibitor or of a PRC2 inhibitor (PRC2Ì). Normalized cell growth is represented in the Iog10 scale, n = 4. P-values were calculated using an unpaired Student’s t-test on day 6 normalized cell counts. For clarity, not all significant differences are displayed in the panel. *P≤0.05. B. Interaction plot representing the day 6 over day 0 ratio (Iog2) of cell counts from serum+LIF +Dox ÷Mock or +PRC2Ì six-day growth assay. * The grey line represents expected results if effects from PRC2 inhibition compared to Mock conditions in TKO H3 cells were additive in TKO H3K4M cells, n = 4. C. Expression-level fold change (log2(RPKM)) for the 130 genes downregulated in TKO H3K4M vs TKO H3 day 0, comparing +PRC2Ì over -PRC2Ì conditions across all cell lines. The dotted line represents log2FC = 1. Each cell line in each condition is represented by three replicates and is grown in serum+LIF +Dox. **D.** Heatmap of row-normalized gene expression (Z-score) for downregulated differentially expressed genes (DEGs (log2FC < -1 and adjusted p-value (t-test) < 0.05); n = 130) in the TKO H3K4M vs TKO H3 comparison in TKO H3K4M grown in -/+PRC2Ì. The mean of two replicates is shown. **A, B, C:** Data are shown as mean ± s.e.m..

### Loss of both 5meC and H3K4me leads to severe defects during pluripotency transitions

Our efforts to examine the dual requirements for 5meC and H3K4me during differentiation were confounded by the severe EB phenotype in TKO cells, independent of whether the H3K4M transgene was expressed. Previous work from our lab showed that TKO ESCs cannot differentiate into the three germ layers but can exit naïve pluripotency (Greenberg et al. 2017; Schulz et al. 2024). Briefly, naïve ESCs cultured in serum-free media supplemented with GSK3β and ERK inhibitors + vitamin C (2i+VitC) exhibit globally low 5meC levels associated with the ground pluripotent state (Nichols and Smith 2009; Habibi et al. 2013; Blaschke et al. 2013); naïve ESCs are then directed to Epiblast-like Cells (EpiLCs) by addition of FGF2 and Activin A (Hayashi et al. 2011). Within two days of EpiLC differentiation, global levels of 5meC are substantially increased, and the *de novo* 5meC program is largely complete by day 4 (Fig. S5A). EpiLCs are still pluripotent but represent the “formative” and “primed” states at day 2 and day 4, respectively (Takahashi et al. 2018). With this system, we reasoned that we would be able to better assess the concerted requirements of 5meC and H3K4me during differentiation.

First, we cultured the four cell lines in naïve conditions. Interestingly, we did not observe any proliferation defect in H3K4M-expressing lines—even in absence of 5meC (Fig. 4A; Fig. S5B). We reasoned that the requirements for H3K4me may be different in this cellular state, as WT naïve ESCs are globally hypomethylated. Next, we differentiated the four lines to EpiLCs (Fig. 4B,C; Fig. S5C). As expected, TKO H3 showed a minimal growth effect (Fig. 4B; Fig. S5C). Moreover, WT H3K4M EpiLCs grew normally despite depleted H3K4me. In contrast, the TKO H3K4M line exhibited a drastic proliferation phenotype, which was exacerbated after day 2 (Fig. 4C). As we did for the serum-grown ESCs, we tested the effect of the DNMT1i on the WT lines, which corroborated the results observed in the TKO background (Fig. S5D,E). However, when we depleted RbBP5 (complete loss of H3K4me), we saw a substantial EpiLC differentiation defect in both WT and TKO lines (Fig. S5F). This prompted us to test whether the H3K4M transgene was being silenced in the WT context, thus masking a possible phenotype. Both flow cytometry and western blot confirmed that the H3K4M transgene was well expressed in WT—albeit slightly lower than TKO in EpiLCs (Fig. S5G,H). Taken together, we showed that proper EpiLC growth requires a threshold of H3K4me. However, absence or reduction of 5meC in an H3K4me-depleted background leads to catastrophic effects.

**Figure 4.**
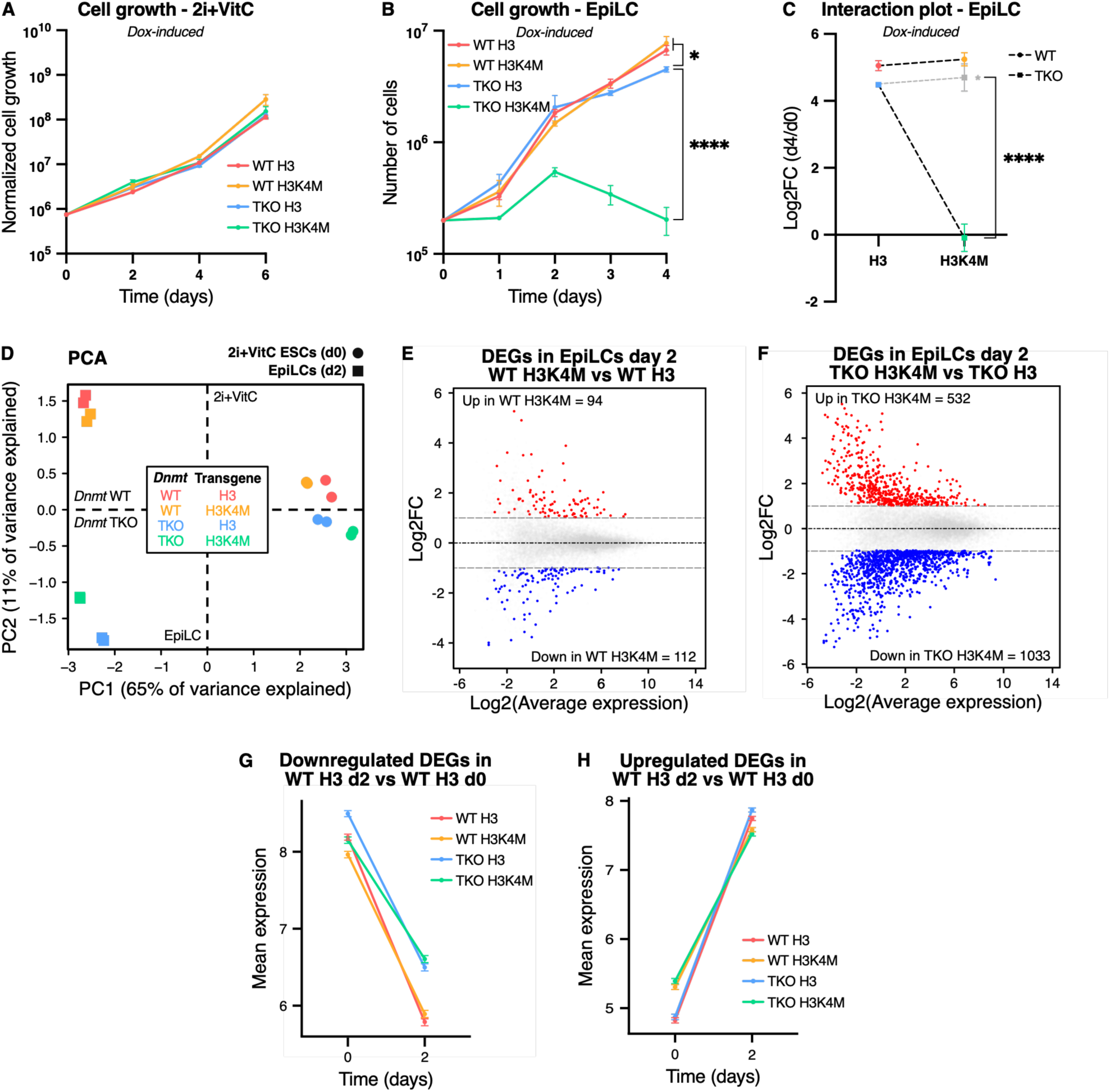
Combined loss of 5meC and H3K4me leads to severe phenotypic and transcriptomic defects during the exit from naive pluripotency. **A.** Six-day growth assay in 2i+VitC +Dox culture conditions. Normalized cell growth is represented in the Iog10 scale, n = 3. B. Four-day growth assay during Epiblast-Like Cells (EpiLC) differentiation (+Dox). Cell numbers are represented in the Iog10 scale, n = 3. **C.** Interaction plot representing the day 4 over day 0 ratio (Iog2) of cell numbers from EpiLC four-day growth assay (+Dox). * The grey line represents expected results if effects from 5meC and H3K4me depletion were additive, n = 3. D. PCA of transcriptomic datasets representing ESCs grown in 2i+VitC (day 0) and EpiLCs (day 2) (+Dox). Each cell line at each time point is represented by two replicates. E, F. MA plots representing differentially expressed genes (DEGs) (|log2FC| > 1 and adjusted p-value (t-test) < 0.05) in EpiLCs day 2 (+Dox) for WT H3K4M vs WT H3 (E) and TKO H3K4M vs TKO H3 (F) comparisons. The average expression is represented as Iog2. Numbers of significantly up- and downregulated genes are indicated. **G, H.** Mean expression levels (log2(RPKM)) for downregulated (G; n = 2456) and upregulated (H; n = 3713) genes in WT H3 during EpiLC differentiation (day 2 vs day 0 comparison) in all cell lines. **A, B, C:** Data are shown as mean ± s.e.m. of replicates from independent experiments. P-values were calculated using an unpaired Student’s t-test on day 4 cell counts (B), or a two-way ANOVA test (C). * P≤0.05, **** P≤0.0001.

Given our previous results, we were curious whether PRC2 inhibition could rescue the TKO H3K4M EpiLC phenotype. TKO H3K4M cells still exhibited major growth impairment, indicating a more important role for PRC2 in EpiLCs than in serum-grown ESCs (Fig. S5I). Nevertheless, we did notice a slight attenuation of the effect of PRC2i on TKO H3K4M compared to TKO H3, suggesting there is a mild phenotypic suppression (Fig. S5J). In sum, the EpiLC system not only bolsters our observations in serum-grown ESCs and EBs but also provides a dynamic system for the interrogation of the interplay of 5meC and H3K4me on genome regulation.

### H3K4M-induced expression defects are exacerbated in TKO H3K4M EpiLCs

To investigate the molecular basis of the TKO H3K4M-associated EpiLC phenotype, we performed a number of genomic analyses on day 0 (naïve pluripotency) and day 2 EpiLCs (formative pluripotency). This timepoint was selected as it appeared to represent an inflection point in the TKO H3K4M phenotype (Fig. 4B), while 5meC levels in WT lines had already been largely established by this stage (Fig. S5A). First, bulk RNA-seq was performed to assess transcriptional dysregulation. Principal component analysis revealed that samples clustered mainly by time point of conversion (Fig. 4D). Cell lines were more similar at day 0, while separation by *Dnmt* background became more pronounced at day 2. Both time points displayed transgene-specific sub-clusters, especially in the case of TKO EpiLCs. Intriguingly, despite no specific growth phenotype in naïve pluripotency (Fig. 4A), pairwise comparisons between H3K4M and H3 lines revealed extensive differential gene expression in both genetic backgrounds, which was exacerbated in the TKO (Fig. S6A,B). Overall, these results highlight the remarkable robustness of the naïve pluripotent state, which tolerates extensive transcriptional rewiring without overt phenotypic consequences.

By day 2 of EpiLC differentiation, gene expression changes were greatly resolved in WT H3K4M compared to WT H3 (total DEGs = 206) (Fig. 4E). Thus, despite transcriptional changes, WT cells were resilient to H3K4me loss in naïve pluripotency and during the transition to formative pluripotency. In contrast, TKO H3K4M EpiLCs displayed massive transcriptional dysregulation (total DEGs = 1565), with the majority (66%) being downregulated relative to TKO H3 (Fig. 4F). In this comparison, GSEA revealed enrichment of developmental terms, including “pattern specification process”, “cell fate commitment” and “embryonic organ development” (Fig. S6C). Similar to day 2 embryoid bodies, roughly half of down-(n = 69) and upregulated (n = 40) DEGs in WT H3K4M were shared with TKO H3K4M, highlighting H3K4M-associated misregulation of these genes (Fig. S6D). Yet, the overall transcriptional effect in TKO H3K4M EpiLCs was much more acute, consistent with the associated phenotypic defects. To investigate in more detail the transition from naïve to formative pluripotency, we compared expression changes in all cell lines. Globally, transcriptome dynamics were preserved among all four conditions: the majority of genes differentially expressed during naïve-to-formative pluripotency transition were shared among the four cell lines (Fig. S6E). In addition, genes typically down- or upregulated in the WT H3 context showed globally similar transcriptional dynamics in the other lines (Fig. 4G,H; Fig. S6E), with minor defects observed in the TKO lines for downregulation, and in the H3KM lines for upregulation. Assessment of naïve pluripotency marker expression indicated that exit of the naïve pluripotent state was achieved in all four conditions by day 2 of conversion (Fig. S6F). Formative markers exhibited misregulation that could appear TKO-specific (e.g., *Fgf5*, *Oct6*), H3K4M-specific (e.g., *Otx2*) or TKO H3K4M-specific (e.g., *Fgfr1*) (Fig. S6G). Moreover, we noted reduced induction of cell junction-associated genes typically expressed during epithelial transitions in TKO H3K4M EpiLCs (Fig. S6H) (Moriwaki et al. 2007; Anderson et al. 2008). Finally, apoptotic factors were specifically upregulated in TKO H3K4M cells at day 2, correlated with the proliferation defects observed in this cell line (Fig. S6I).

Having established that the TKO H3K4M line exhibited the most misregulated transcriptome, we next sought to determine how the combined loss of H3K4me and 5meC affects chromatin states at regulatory elements, and how this might contribute to gene expression changes.

### H3K4me1 loss drives distinct repressive states in 5meC-present or -absent contexts

We first set out to determine the impact of H3K4M expression on enhancers, which are generally correlated with the presence of H3K4me1 (Barral and Déjardin 2023). During cell-fate transitions, the enhancer landscape is extensively remodelled to support new transcriptional programs (Buecker and Wysocka 2012; Whyte et al. 2012). To analyze enhancers in our cell lines, we used a recently established catalog of candidate enhancers in naïve ESCs and EpiLCs (Morales-Franco et al. 2026). Briefly, candidate enhancers positive for H3K4me1 and H3K27 acetylation (H3K27ac) and displaying accessibility (assessed by ATAC-seq) were defined as active; accessible enhancers enriched for H3K4me1 but lacking H3K27ac signal were classified as primed; non-promoter regions marked only by H3K4me1 were categorized as inaccessible putative enhancers (Fig. 5A).

**Figure 5.**
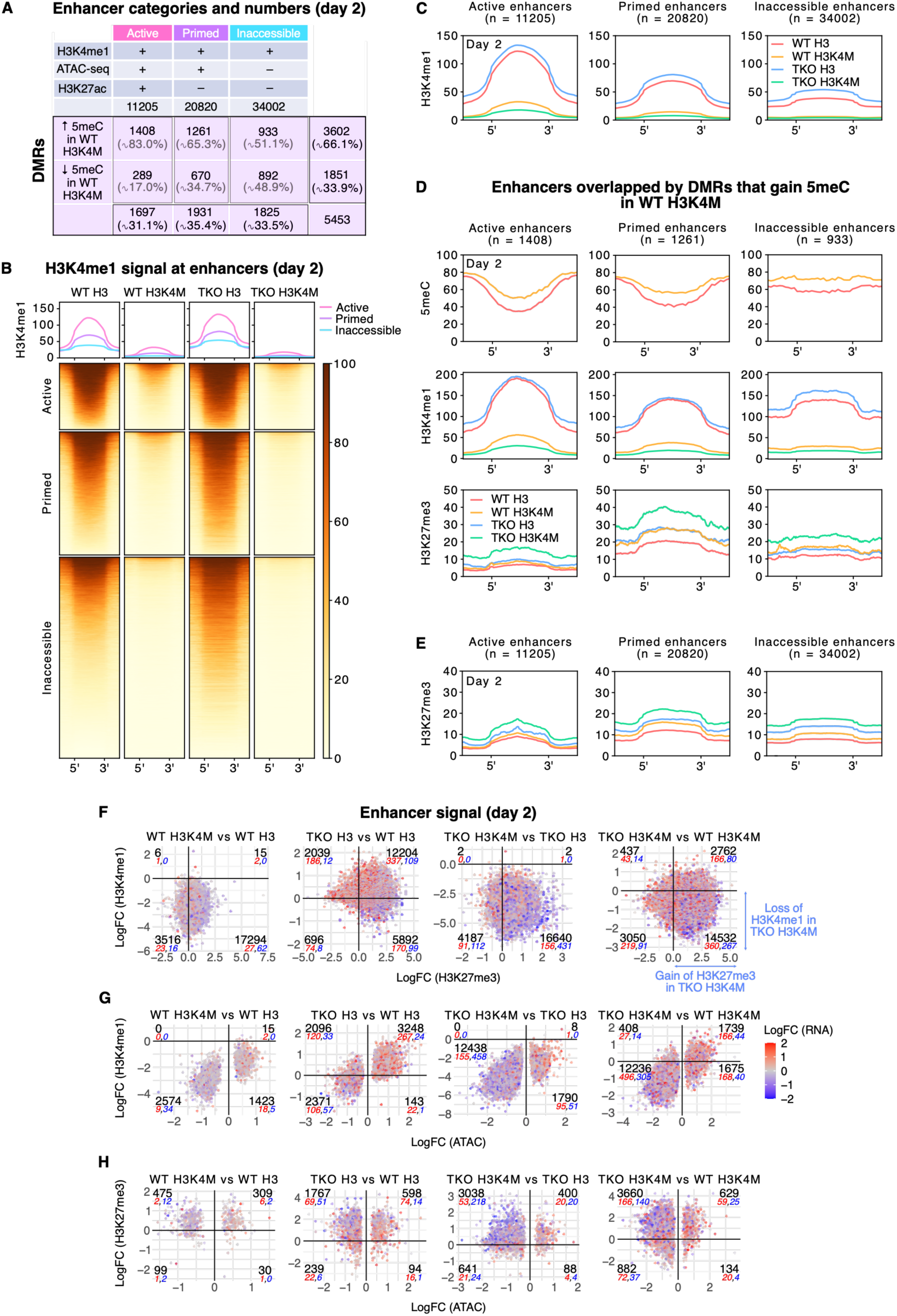
H3K4me1 depletion at enhancers disrupts other chromatin features. **A.** Table showing numbers and associated chromatin features for each category of enhancers (active, primed, inaccessible) as defined in (Morales-Franco et al. 2026) (top), together with number of differentially methylated regions (DMRs) overlapping enhancers and gaining or losing 5meC in WT H3K4M vs WT H3 (bottom). Percentages in grey represent the proportion among all DMRs identified within each enhancer category. Percentages in black (bottom) represent the proportion of each enhancer category among all DMRs, and percentages in black (right) represent the proportion of enhancer DMRs either gaining or losing 5meC in WT H3K4M. All numbers are from EpiLCs (day 2). **B, C.** Heatmap **(B)** and profile plots **(C)** of normalized H3K4me1 signal at enhancers and 1 kb upstream and downstream in EpiLCs (day 2). Enhancers are clustered by category. The mean of two replicates is shown. D. Profile plots of 5meC, normalized H3K4me1 and normalized H3K27me3 signals at enhancer DMRs that gain 5meC in WT H3K4M vs WT H3, and 1 kb upstream and downstream. Enhancers are clustered by category and signal is shown for all cell lines in EpiLCs (day 2). The mean of two replicates is shown. E. Profile plots of normalized H3K27me3 signal at enhancers clustered by category in EpiLCs (day 2). The mean of two replicates is shown. F. Scatterplots showing the fold-change of H3K27me3 signal (Iog2) vs the fold-change of H3K4me1 signal (Iog2) at enhancers in the different comparisons, colored by the fold-change of enhancer-associated gene expression (log2(RPKM)) in EpiLCs (day 2). **G, H.** Scatterplots showing the fold-change of ATAC signal (Iog2) vs the fold-change of H3K4me1 **(G)** and H3K27me3 (H) signal (Iog2) at enhancers in the different comparisons, colored by the fold-change of enhancer-associated gene expression (log2(RPKM)) in EpiLCs (day 2). Only enhancers with statistically-significant differential accessibility (FDR < 0.05) are depicted. For F-H, negative values indicate signal loss in the first cell line of each comparison, whereas positive values indicate signal gain in the first cell line. The number of enhancers (black) as well as the number of associated significantly upregulated (red) or downregulated (blue) DEGs are indicated in each quadrant.

H3K4me1 CUT&Tag revealed extensive depletion of the mark across all enhancer classes in both H3K4M lines at both time points, with the strongest loss in TKO H3K4M (Fig. 5B,C; Fig. S7A,B). Notably, TKO H3 EpiLCs displayed higher H3K4me1 levels than WT H3 EpiLCs, consistent with our MS data in serum-grown ESCs (Fig. S1A).

In order to gain insights into a potential regulatory role 5meC plays at enhancer elements, we performed Whole-Genome Bisulfite Sequencing (WGBS) in WT H3 and H3K4M EpiLCs. We used MethyLasso to define differentially methylated regions (DMRs) genome-wide (Balaramane et al. 2024), and intersected them with our enhancer regions. We observed enhancers exhibiting a gain of 5meC (66%) or a loss (34%) in WT H3K4M cells (Fig. 5A), associated with distinct chromatin states. Enhancers losing the 5meC mark were predominantly inaccessible (892/1851; 48.2%) and primed (670/1851; 36.2%), and generally harbored elevated 5meC and low H3K4me1 levels in WT H3 EpiLCs (Fig. 5A; Fig. S7C). In contrast, the enhancers gaining 5meC in WT H3K4M EpiLCs were mainly active (1408/3602; 39.1%) and primed (1261/3602; 35%) and, in WT H3, exhibited higher H3K4me1 signal as well as lower baseline 5meC compared to enhancers losing 5meC in WT H3K4M (Fig. 5A,D). While the skewed gain of 5meC towards active and primed enhancers may have been incidental, based on our phenotypic assays, we were motivated to determine if the methyl mark might play a compensatory role.

Given our observations that a PRC2 inhibitor can suppress TKO H3K4M cellular phenotypes in serum ESCs, we went on to assess H3K27me3 levels at enhancer elements in day 2 EpiLCs, when genome-wide 5meC levels are high in WT. Globally, H3K27me3 signal increased in all enhancer categories in both H3K4M lines compared to respective H3 controls, with TKO H3K4M EpiLCs clearly showing the strongest enrichment (Fig. 5E,F). This was particularly striking at active enhancers, where gain of the repressive mark was substantial in TKO H3K4M compared to the TKO H3 line, while increase in the WT H3K4M line was minimal (Fig. 5E). When focusing on active enhancers gaining 5meC, our analysis was particularly revealing: in WT H3K4M, those enhancers were not particularly enriched for H3K27me3, but by contrast, this same set of regulatory elements displayed clear accumulation of H3K27me3 in TKO H3K4M. This suggests that, in WT H3K4M, ectopic gain of 5meC at a subset of enhancers may contribute to limiting H3K27me3 deposition (Fig. 5D). Overall, H3K4me1 loss at enhancers leaves them susceptible to H3K27me3 accumulation, especially in the primed and inaccessible class (Fig. 5E). However, this effect is exaggerated in the TKO (Fig. 5E,F; TKO H3K4M vs WT H3K4M comparison, bottom right quadrant), supporting a combined role for H3K4me1 and 5meC in restricting the PRC2-deposited mark at enhancers.

In line with a transition to a more repressed state, we also observed that loss of H3K4me1 at enhancers was generally correlated with decreased chromatin accessibility, as shown by ATAC-seq (Fig. 5G). Active and primed rather than inaccessible enhancers displayed the most accessibility loss across all comparisons (Fig. S7D). Once again, the effect was most pronounced in the TKO H3K4M line. In addition, despite the general accumulation of H3K27me3 at enhancers in WT H3K4M when compared to WT H3 (Fig. 5E), limited changes in accessibility were observed, while the additional absence of 5meC appeared to exacerbate accessibility loss (Fig. 5H). A subset of enhancers displaying a gain of 5meC in WT H3K4M cells was predominantly less accessible than in WT H3 (Fig. S7E)—a feature consistent with the classical view of 5meC as repressive. Yet, in TKO H3K4M EpiLCs, accessibility of an even greater proportion of this set of enhancers was decreased. Hence, H3K4me1 and 5meC appear to play semi-redundant roles in creating a chromatin environment amenable for transcription factor binding.

Finally, we reasoned that the stark impact on enhancer state would be reflected in decreased gene expression. To assess this, we used Genomic Regions Enrichment of Annotations Tool (GREAT) results obtained from (Morales-Franco et al. 2026) to link enhancers to potential target genes (McLean et al. 2010). We integrated our enhancer results on chromatin marks and accessibility with gene expression changes in day 2 EpiLCs (Fig. 5F-H). Downregulated genes in TKO H3K4M comparisons were frequently associated with enhancers displaying combined loss of H3K4me1 and gain of H3K27me3, as well as loss of accessibility (numbers of significantly downregulated genes are represented in blue). Similarly, enhancers that gained H3K27me3 were preferentially associated with downregulated genes in the TKO H3K4M comparisons (TKO H3K4M vs TKO H3 and TKO H3K4M vs WT H3K4M), whereas this association was weaker in WT or H3-only comparisons (Fig. S7F). Therefore, although these observations are correlative, they suggest that the disruption of the enhancer landscape resulting from absence of 5meC and reduction of H3K4me1 may contribute to the alteration of gene expression programs, potentially through the accumulation of H3K27me3 and reduced accessibility at a sizeable proportion of enhancers. These observations are exemplified by a putative enhancer predicted to regulate the *Tle3* gene (Fig. S7G), which is downregulated in TKO H3K4M EpiLCs and whose regulation will be discussed in more detail below.

Altogether, these results revealed distinct chromatin responses in WT and TKO backgrounds. The gain of H3K4me1 in TKO H3, and the increase in 5meC at active enhancers in WT H3K4M both point to compensatory mechanisms. The pronounced chromatin alterations observed in TKO H3K4M EpiLCs, characterized by the highest H3K27me3 enrichment, least chromatin accessibility and greatest association with gene expression downregulation, motivated us to explore the impact at gene promoters.

### Promoter H3K4me3 and associated gene expression is most affected in the TKO H3K4M context

To assess the requirement of H3K4me3 and 5meC at promoter regions, we first subgrouped promoters into CGI and non-CGI classes. We reasoned that constitutively unmethylated CGI promoters might exhibit different regulatory properties than the non-CGI class, which display globally higher 5meC levels (Stadler et al. 2011). Secondly, we subgrouped each class into bivalent and non-bivalent promoters using our CUT&Tag datasets. Bivalent promoters are often associated with developmental genes, and are defined by the dual presence of H3K4me3 and H3K27me3 peaks (Azuara et al. 2006; Bernstein et al. 2006). We noticed a limited H3K4me3 signal across all cell lines at non-CGI promoters at both time points, while CGI promoters displayed higher and more interpretable levels (Fig. 6A,B; Fig. S8A,B). At day 0, we observed a moderate decrease in H3K4me3 in the WT H3K4M line compared to WT H3, and virtually no effect in the TKO H3K4M (Fig. S8A,B). This result was curious, given the greater transcriptional misregulation we observed at day 0 TKO H3K4M (Fig. S6B). However, we observed an inverted effect on day 2: while the signal in WT H3K4M remained reduced compared to WT H3, TKO H3K4M EpiLCs displayed the lowest H3K4me3 signal at all promoter categories (Fig. 6A,B). Despite a substantial effect detected in day 2 H3K4M lines— especially in TKO H3K4M—these results stood in stark contrast to the more drastic global effect observed for H3K4me1 in the same lines.

**Figure 6.**
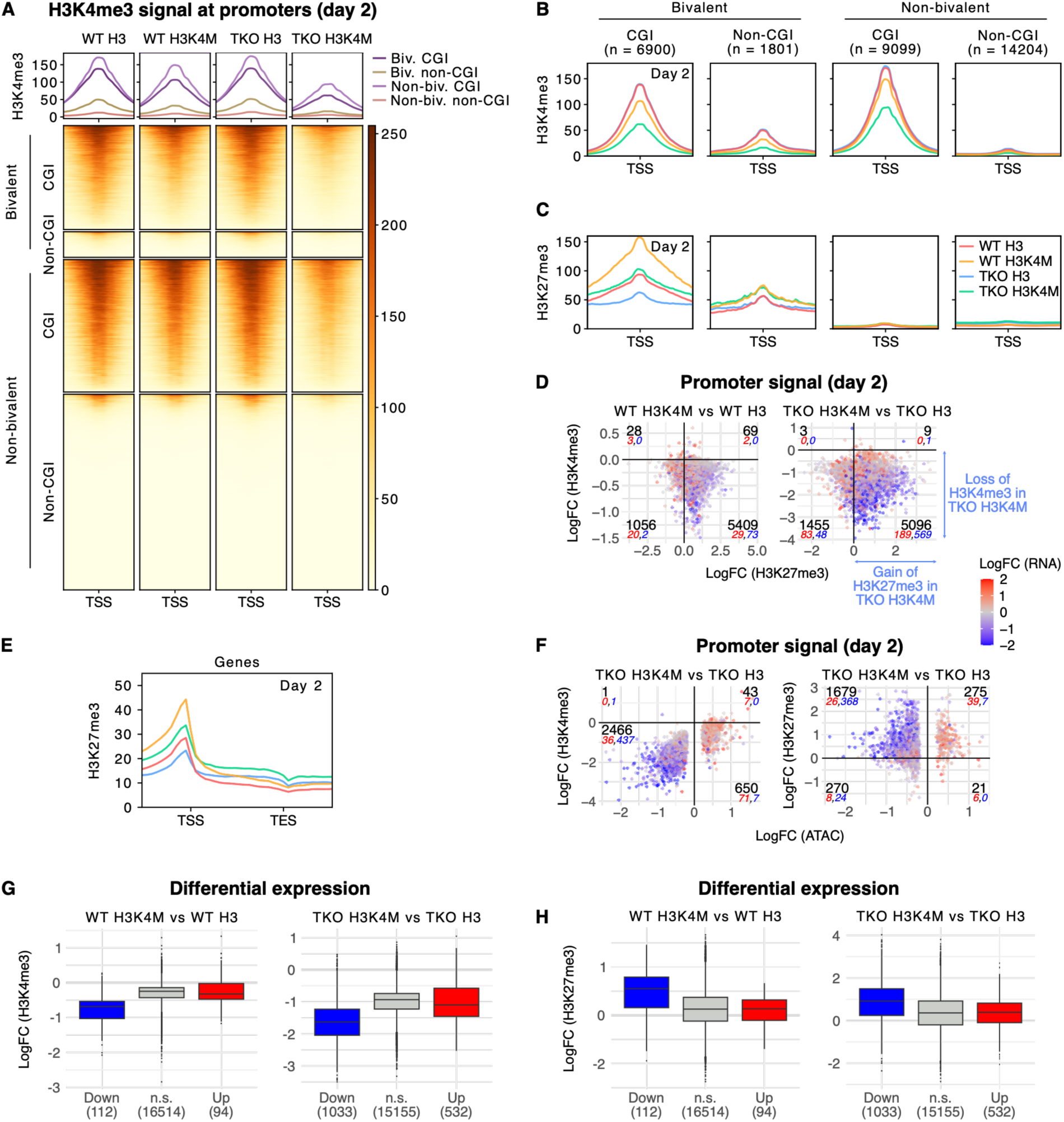
H3K4me3 depletion at gene promoters results in gain of H3K27me3. **A, B.** Heatmap **(A)** and profile plots **(B)** of normalized H3K4me3 signal at bivalent and non-bivalent promoters, splitted into CGI and non-CGI promoters (± 2 kb around the TSS) in EpiLCs (day 2). The mean of two replicates is shown. C. Profile plot of normalized H3K27me3 signal at bivalent and non-bivalent promoters, splitted into CGI and non-CGI promoters (± 2 kb around the TSS) in EpiLCs (day 2). The mean of two replicates is shown. **D.** Scatterplots showing the fold-change of H3K27me3 signal (Iog2) vs the fold-change of H3K4me3 signal (Iog2) at promoters for H3K4M vs H3 comparisons, colored by the fold-change of associated gene expression (log2(RPKM)) in EpiLCs (day 2). Negative values indicate signal loss in the first cell line of each comparison, whereas positive values indicate signal gain in the first cell line. The number of promoters (black) as well as the number of associated significantly upregulated (red) or downregulated (blue) DEGs are indicated in each quadrant. E. Profile plot of normalized H3K27me3 signal from the Transcription Start Site (TSS) to the Transcription End Site (TES) in EpiLCs (day 2) protein coding genes, as well as 1 kb upstream and downstream. For visualization, data are shown for genes located on the + strand. The mean of two replicates is shown. **F.** Scatterplots showing the fold-change of ATAC signal (Iog2) vs the fold-change of H3K4me3 (left) or H3K27me3 (right) signal (Iog2) at promoters for TKO H3K4M vs TKO H3 comparison, colored by the fold-change of associated gene expression (log2(RPKM)) in EpiLCs (day 2). Only promoters with statistically-significant differential accessibility (FDR < 0.05) are depicted. Negative values indicate signal loss in TKO H3K4M cells, whereas positive values indicate signal gain in this cell line. The number of promoters (black) as well as the number of associated significantly upregulated (red) or downregulated (blue) DEGs are indicated in each quadrant. **G, H.** Boxplots showing fold-change of H3K4me3 signal (Iog2) **(G)** and H3K27me3 signal **(H)** at promoters of downregulated (Down), non-significant (n.s.) or upregulated (Up) genes (|log2FC| > 1 and adjusted p-value (t-test) < 0.05) in the H3K4M vs H3 comparisons in EpiLCs (day 2).

We then went on to interrogate how 5meC at gene promoters is impacted and how this might influence their regulation. Similar to enhancers, DMRs both gaining (2305; 7.2% of total gene promoters) and losing 5meC (2507; 7.8% of total gene promoters) were identified at gene promoters in WT H3K4M EpiLCs, indicating fairly widespread 5meC misregulation at regulatory elements (Fig. S8C,D). At non-CGI promoters losing 5meC, we observed a limited H3K4me3 signal associated with high 5meC levels in both WT lines. In contrast, those gaining 5meC displayed higher H3K4me3 levels than those losing 5meC, with decreased H3K4me3 levels in both H3K4M lines (Fig. S8C). Pronounced H3K4me3 depletion in H3K4M lines was observed at CGI promoters overlapping DMRs showing either gains or losses of 5meC, although the associated changes in 5meC levels were generally modest.

Next, we wanted to assess how H3K27me3 was impacted at promoters (Fig. 6C; Fig. S8E). Similarly to the H3K4me3 signal, H3K27me3 levels at day 2 were lower at bivalent non-CGI promoters compared to CGI ones, consistent with PRC2 targeting unmethylated CpG-rich promoters (Blackledge and Klose 2021) (Fig. 6C). While a minimal effect was observed on day 0 at both CGI and non-CGI promoters (Fig. S8E), H3K27me3 levels were strongly increased in both H3K4me-depleted lines in day 2 compared to respective controls. Notably, this increase was almost exclusively confined to bivalent promoters associated with a loss of H3K4me3 enrichment (Fig. 6C,D). We observed a higher amplitude of the normalized H3K27me3 signal in WT H3K4M compared to TKO H3K4M at bivalent CGI promoters in EpiLCs, matching differences already visible between H3 lines, while bivalent non-CGI promoters displayed a similar increase of the signal between both H3K4M lines (Fig. 6C). Therefore, bivalent promoters appear particularly sensitive to gain of H3K27me3 following loss of H3K4me3 in H3K4M lines, as previously reported (Yagi et al. 2025). H3K4me3 and H3K27me3 occur on asymmetric histones in bivalent nucleosomes (Voigt et al. 2012), and the increased H3K27me3 signal in H3K4M lines likely reflects PRC2 activity on both histone tails. This also indicates that PRC2 is not nucleating on *de novo* targets genome-wide when H3K4me3 is reduced.

We were curious if 5meC dynamics in the H3K4M lines could impact the H3K27me3 landscape. Interestingly, promoters overlapping DMRs exhibited similar changes in H3K27me3 levels, particularly at CGI promoters (Fig. S8C,D). The concomitant gain of 5meC and H3K27me3 in WT H3K4M cells is notable, given that these marks are typically antagonistic at CpG-rich regions (Statham et al. 2012; Brinkman et al. 2012). One possible explanation is that 5meC levels at these DMRs remained below the threshold required to counteract H3K27me3 deposition. Interestingly, bivalent non-CGI promoters were more susceptible to gain of 5meC (392/1801; 21.8%) than other types of promoters (Fig. S8D), suggesting that 5meC accumulation may partially compensate for H3K27me3 deposition in WT H3K4M, akin to what we observed at a subset of active enhancers.

Nevertheless, among all promoter classes, a portion exhibited increased gain of H3K27me3 in TKO H3K4M relative to other cell lines (Fig. S8F; TKO H3K4M vs WT H3K4M comparison, bottom right quadrant). In addition, in the TKO lines, we observed increased spreading of the H3K27me3 mark over gene bodies, with the highest level in TKO H3K4M (Fig. 6E). To summarize, it appears H3K4me3 restrains H3K27me3 establishment at the TSS of bivalent promoters, while 5meC over the gene body prevents the spreading of the mark (King et al. 2016; Richard Albert et al. 2025; Yagi et al. 2025). Again, this suggests an intriguing cooperative role of H3K4me and 5meC.

Despite the chromatin phenotype observed at WT H3K4M promoters in day 2 EpiLCs, very little differential accessibility was observed, and was not associated with H3K4me3 loss nor H3K27me3 gain (Fig. S8G,H). Additionally, the loss of promoter accessibility in TKO H3 compared to WT H3 was not entirely correlated with H3K27me3 changes (Fig. S8H). In contrast, in the TKO H3K4M line, the gain of promoter H3K27me3 was correlated with loss of accessibility in comparison to TKO H3 (Fig. 6F). However, the H3K4me3 signal loss is most acute at gene promoters in TKO H3K4M, and may be the main driver of reduced accessibility (Fig. 6F; Fig. S8G; TKO H3K4M vs WT H3K4M comparison), as previously suggested (Barsoum et al. 2022).

Finally, we wondered how gene expression was affected following H3K4me3 loss and 5meC and/or H3K27me3 gain at gene promoters, as well as accessibility changes. Importantly, 37 genes showing gain of 5meC at CGI and non-CGI gene promoters were significantly downregulated in WT H3K4M, thus representing 33% of downregulated DEGs in WT H3K4M vs WT H3 (Fig. 4E; Fig. S8D). Therefore, part of the significant gene expression downregulation in WT H3K4M could be explained by the gain of repressive 5meC at gene promoters. However, this represents only a fraction (1.6%) of the total 2305 promoters gaining 5meC. Importantly, gene expression downregulation appeared to be highly correlated with loss of promoter accessibility when associated with H3K4me3 loss or H3K27me3 gain, particularly in the TKO H3K4M line (Fig. 6F; Fig. S8G,H). Similar to H3K4me1, H3K4me3 loss was not entirely predictive of downregulation of gene expression in H3K4M lines (Fig. 6D; Fig. S8F). The relationship between H3K27me3 and gene repression appeared paradoxical: promoter H3K27me3 was more prominently gained in WT H3K4M than in TKO H3K4M at bivalent CGI promoters, yet transcriptional repression was markedly stronger in TKO H3K4M (Fig. 4E,F; Fig. 6C,D). More generally, across all conditions, significantly downregulated genes are characterized by a stronger loss of H3K4me3 and a stronger gain of H3K27me3 (Fig. 6G,H; Fig. S8I,J). In sum, integrating all data sets suggests that H3K4me3 loss or H3K27me3 gain alone do not predict changes in gene expression, but downregulated genes exhibit both these features, and H3K27me3 spreading over gene bodies in TKO H3K4M may contribute to an increased repressive state.

### Epigenome editing demonstrates chromatin dynamics at selected promoters

Our data point to a three-way interplay between H3K4me, 5meC and H3K27me3 for proper gene regulation. However, the TKO mutations and the inducible H3K4M transgene both lead to global misregulation of the epigenome and transcriptome, which could result in secondary effects that may confound our interpretations at individual loci. To functionally test the role of the chromatin marks, we employed two epigenome editing strategies. Briefly, we used a constitutively expressed catalytically inactive Cas9 (dCas9) fused to five copies of the GCN4 epitope tag (SunTag) (Tanenbaum et al. 2014). The SunTag is recognized by a single-chain variable fragment (scFv) fused to the catalytic domain of a chromatin-modifying enzyme. For our first strategy, we used the catalytic domain of TET1, which leads to DNA demethylation (Morita et al. 2016; Horii et al. 2022; Richard Albert et al. 2025). We expressed dCas9-SunTag/scFv-GFP-TET1 (dCas9-ST/TET1) in the WT H3K4M line, allowing us to toggle global H3K4me levels by adding Dox to the medium (Fig. 7A, left). As a control, we used dCas9-ST/TET1 with scrambled guides. For our second strategy, we used the catalytic domain of the H3K4 demethylase KDM5B (dCas9-ST/KDM5B) (Xhabija and Kidder 2019), which we expressed in WT and TKO H3 backgrounds (Fig. 7A, right). As a control, we used a catalytically inactive version of KDM5B (KDM5B CI). To summarize, with these tools, we could locally demethylate DNA in a globally H3K4-hypomethylated background, or locally demethylate H3K4 in a globally DNA-hypomethylated background.

**Figure 7.**
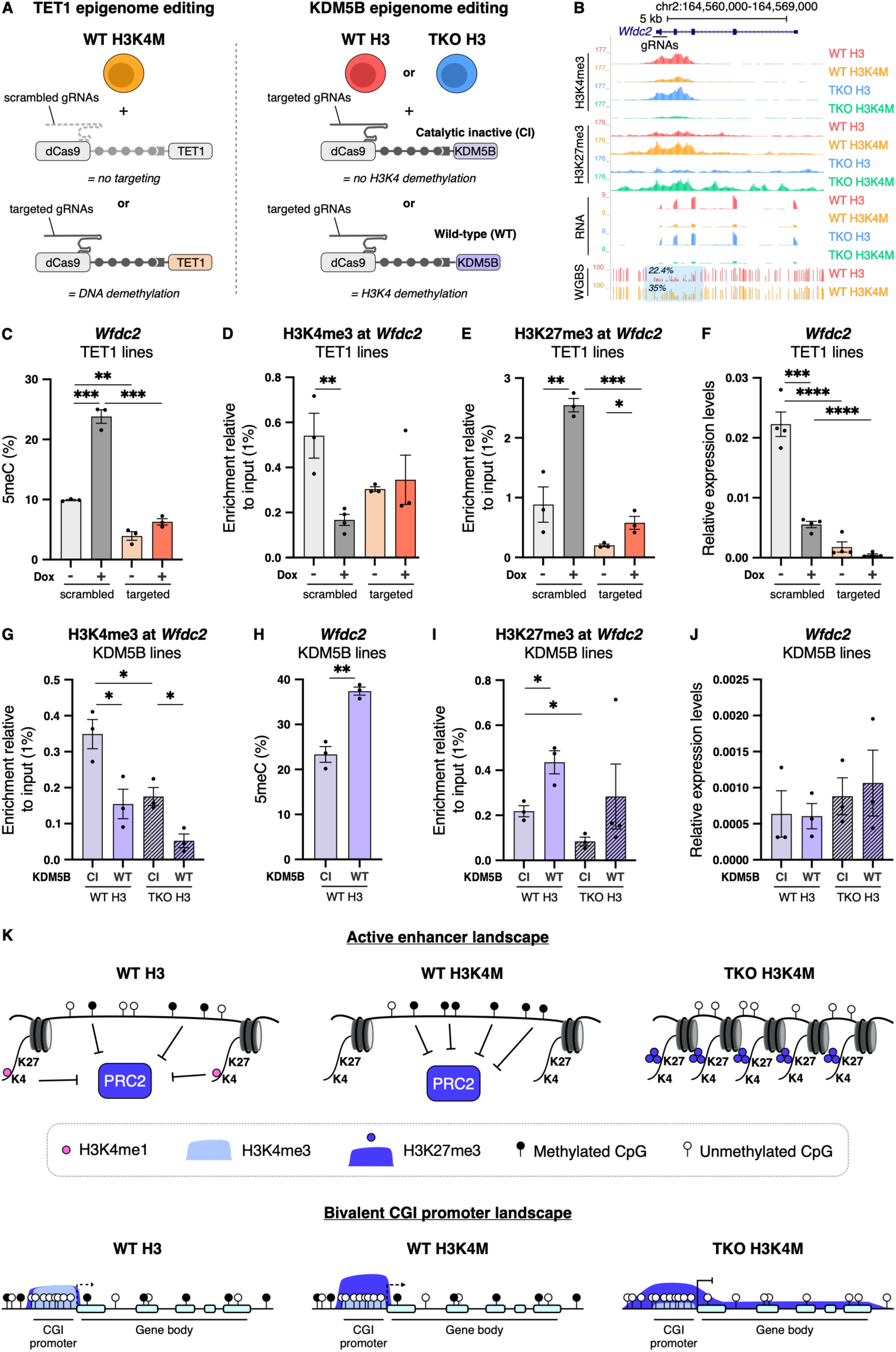
Targeted depletion of 5meC or H3K4me3 results in local chromatin changes at *Wfdc2* promoter. **A.** Schematic of the epigenome editing targeted strategy at candidate promoters. Site-specific DNA demethylation is carried out by constitutive expression of dead Cas9 (dCas9)-SunTag/TET1 catalytic domain (CD) and gRNAs targeted to candidate promoters in the WT H3K4M line. Cells constitutively expressing scrambled gRNAs (untargeted) are used as a control (left). Site-specific H3K4 demethylation is carried out by constitutive expression of dCas9-SunTag/KDM5B WT and gRNAs targeted to candidate promoters in WT H3 and TKO H3 lines. Control lines expressed a targeted catalytic-inactive (Cl) version of the KDM5B construct (right). **B.** UCSC Genome Browser screenshot of the *Wfdc2* locus in all cell lines in EpiLCs (day 2). The region targeted by gRNAs is indicated. From top to bottom, normalized H3K4me3 and H3K27me3 CUT&Tag signals, RNA-seq signal, and 5meC levels in WT lines are shown. Average 5meC levels over the highlighted region are indicated in WT lines. **C.** 5meC levels assessed by bisulfite-pyrosequencing at the *Wfdc2* promoter in TET1 epigenome editing lines. Assay was performed in EpiLCs (day 2) with uninduced or induced H3K4M expression (-/+ Dox). Each data point represents the average 5meC levels over the pyrosequencing amplicon (composed of eleven CpGs) for each replicate. Replicates and their mean are shown, n = 3. D, E. H3K4me3 (D) and H3K27me3 (E) levels assessed by ChlP-qPCR at the *Wfdc2* promoter in TET1 epigenome editing lines in EpiLCs (day 2) with uninduced or induced H3K4M expression (-/+ Dox). Enrichment is calculated relative to input (1%). Replicates and their mean are shown, n = 3; except for H3K4me3 in scrambled +Dox, n = 4. F. *Wfdc2* expression assessed by RT-qPCR in TET1 epigenome editing lines in EpiLCs (day 2) with uninduced or induced H3K4M expression (-/+ Dox). Relative expression levels are normalized to the average Ct of the two housekeeping genes *Rrm2* and *RpIpO* using the ΔCt method, n = 4. G. H3K4me3 levels assessed by ChlP-qPCR at the *Wfdc2* promoter in KDM5B epigenome editing lines in EpiLCs (day 2). Enrichment is calculated relative to input (1%). Replicates and their mean are shown, n = 3. H. 5meC levels assessed by bisulfite pyrosequencing at the *Wfdc2* promoter in KDM5B epigenome editing lines in WT EpiLCs (day 2). Each data point represents the average 5meC levels over the pyrosequencing amplicon (composed of eleven CpGs) for each replicate. Replicates and their mean are shown, n = 3.1. H3K27me3 levels assessed by ChlP-qPCR at the *Wfdc2* promoter in KDM5B epigenome editing lines in EpiLCs (day 2). Enrichment is calculated relative to input (1%). Replicates and their mean are shown, n = 3; except TKO H3 KDM5B WT, n = 4. **J.** *Wfdc2* expression assessed by RT-qPCR in KDM5B epigenome editing lines in EpiLCs (day 2). Relative expression levels are normalized to the average Ct of the two housekeeping genes *Rrm2* and *RpIpO* using the ΔCt method, n = 3. **K.** Proposed models for the cooperative role of 5meC and H3K4me at active enhancers and bivalent CGI promoters. At active enhancers (top), the presence of H3K4me1 and DNA methylation (black lollipops) in WT cells (left) contributes to maintaining an active enhancer state and may impede the establishment of Polycomb-mediated silencing via the PRC2 complex. In WT H3K4M cells (middle), loss of H3K4me1 leads to gain of 5meC at a subset of active enhancers, maintaining protection from PRC2-mediated silencing. In *Dnmt* TKO H3K4M cells (right), the absence of DNA methylation and depletion of H3K4me1 allows PRC2-mediated deposition of H3K27me3 and associated enhancer silencing. At bivalent CGI gene promoters (bottom), in WT cells (left), the presence of H3K4me3 associated with DNA methylation across gene bodies and at CGI shores prevent the accumulation and spreading of H3K27me3. In WT H3K4M cells (middle), reduced H3K4me3 at bivalent promoters is associated with a substantial gain of H3K27me3 at these regulatory elements. Despite these epigenetic changes, transcription is generally maintained, suggesting that the gain of H3K27me3 alone is insufficient to induce global transcriptional repression. In TKO H3K4M cells (right), the combined absence of 5meC and depletion of H3K4me3 allows accumulation and spreading of H3K27me3, which contributes to transcriptional repression. **C-J:** Data are shown as mean ± s.e.m. of replicates from independent experiments. P-values were calculated using an unpaired Student’s t-test **(C-l). \*** P≤0.05, ** P≤0.01, *** P≤0.001, **** P≤0.0001.

We opted to target promoters, rather than enhancers, reasoning that we would be able to more directly assess the effect on gene expression. We chose loci based on the following criteria observed in day 2 EpiLCs: 1) decrease of H3K4me3 and increase of H3K27me3 in the H3K4M lines, with spreading of H3K27me3 into the gene body in the TKO H3K4M; and 2) decreased gene expression in the H3K4M lines, with the strongest effect in the TKO H3K4M. We chose two bivalent genes, one with a non-CGI promoter, *Wfdc2* (Fig. 7B), and one with a CGI promoter, *Tle3* (Fig. S9A). In addition, the *Wdfc2* promoter overlapped a DMR gaining 5meC in WT H3K4M. While the*Tle3* promoter did not overlap a DMR, we did note a minor gain of 5meC at the *Tle3* promoter region (average 5meC levels in WT H3: 11.2%; in WT H3K4M: 15.6%).

Using our first strategy, we were able to demonstrate dCas9-ST/TET1-mediated demethylation of *Wfdc2* in both the uninduced and induced WT H3K4M line (Fig. 7C). The *Tle3* CGI promoter was already hypomethylated, thus was only minimally impacted by the dCas9-ST/TET1 (Fig. S9B). Next, we performed both H3K4me3 and H3K27me3 ChIP-qPCR to assess the effect of TET1 editing on histone marks (Fig. 7D,E; Fig. S9C-F). At the *Wfdc2* locus, a decrease in H3K4me3 was observed following H3K4M induction, but this was not further exacerbated with the dCas9-ST/TET1 editing (Fig. 7D). Moreover, the increase of H3K27me3 associated with H3K4M expression was impeded by the presence of the epigenome editing machinery (Fig. 7E). Gene expression assessed by RT-qPCR showed that the dCas9-ST/TET1 targeting led to transcriptional interference, thus making the DNA demethylation effect difficult to parse (Fig. 7F). We only observed a minor effect on H3K4me3 at the *Tle3* locus (Fig. S9E), and once again presence of the epigenome editing machinery appeared to negatively affect the increase in H3K27me3 signal (Fig. S9F). Interestingly, we did observe a specific, if statistically insignificant, effect of expression when dCas9-ST/TET1 was targeted in tandem with H3K4M induction (Fig. S9G). This might indicate a H3K27me3 spreading effect that reinforces silencing that occurred in TKO H3K4M (Fig. S9A). Despite efficient DNA demethylation at *Wfdc2* and a notable RT-qPCR response at *Tle3*, the broader functional impact of dCas9-ST/TET1 remains to be fully defined. This motivated our shift to the second strategy.

After establishing the dCas9-ST/KDM5B lines and respective controls, we first assessed the efficacy of the construct by H3K4me3 ChIP-qPCR (Fig. 7G; Fig. S9H,I). The editing did not appear to function as hoped at the *Tle3* locus. While we did note a slight increase in H3K27me3 signal in the TKO line harboring dCas9-ST/KDM5B relative to KDM5B CI (Fig. S9J) concomitant with a decrease in expression (Fig. S9K), the lack of an effect at H3K4me3 makes this difficult to interpret. In contrast, we were encouraged that dCas9-ST/KDM5B led to strong reduction in H3K4me3 at *Wfdc2* in both WT and TKO (Fig. 7G). This reduction was correlated with a gain in both 5meC (Fig. 7H) and H3K27me3 (Fig. 7I, Fig. S9L). Hence, we locally recreated the precise chromatin dynamics that we observed from the genome-wide data (Fig. 7B). However, we did not observe a decrease in expression in the dCas9-ST/KDM5B line compared to KDM5B CI (Fig. 7J), but it should be noted that simply the presence of the dCas9 machinery at the *Wfdc2* promoter leads to a decrease of expression by an order of magnitude, independently of which epigenetic modifier it is recruiting (Fig. 7F); perhaps it is unsurprising that no further reduction occurred in dCas9-ST/KDM5B lines, despite the effect on chromatin. Nevertheless, our analyses at the *Wfdc2* locus demonstrate that modifying the H3K4me3 state has an effect on both 5meC and H3K27me3, indicating the protective role it plays at promoter elements.

## Discussion

Employing inducible H3K4M mouse ESC lines, we investigated the extent to which H3K4me protects regulatory elements from DNA methylation during cell-fate transitions—a process that requires establishing and maintaining specific gene expression programs. In our system, H3K4me depletion resulted in rather limited global 5meC deposition at regulatory elements during the transition from naïve pluripotent ESCs to formative EpiLCs. Instead, we observed accumulation of the repressive histone modification H3K27me3 at both enhancers and bivalent promoters, which was further enhanced when H3K4me depletion was combined with the absence of 5meC. Importantly, H3K27me3 gain in the latter context was associated with reduced regulatory element accessibility, increased transcriptional dysregulation, and exacerbated cellular phenotypic defects in pluripotent and differentiating cells. Together, our results revealed an unexpected functional interplay between two pathways generally considered mutually exclusive in limiting PRC2-mediated repression during pluripotency and cell-fate transitions.

Our data have allowed us to propose two related, but distinct, models of regulatory element regulation. At enhancers, TKO H3K4M EpiLCs revealed a requirement for the presence of H3K4me1 and 5meC. In the absence of both marks, enhancers globally displayed pronounced loss of accessibility concomitant with H3K27me3 accumulation, consistent with loss of activity. Importantly, these changes at enhancers were associated with a tendency toward downregulation of associated gene expression. Together, these findings demonstrate that H3K4me1 and 5meC can function combinatorially to protect enhancers from PRC2-mediated repression (Fig. 7K, top). In agreement with such a model, both marks have been reported to limit PRC2 occupancy (Schmitges et al. 2011; Brinkman et al. 2012; Statham et al. 2012; Tvardovskiy et al. 2024). Moreover, the ectopic gain of 5meC at a subset of active and primed enhancers in WT H3K4M cells, together with the specific accumulation of H3K27me3 at these elements in TKO H3K4M cells, raises the possibility that 5meC can partially compensate for the loss of H3K4me1 at enhancers. Conversely, in TKO H3 cells lacking DNA methylation, increased H3K4me1 levels at enhancers suggest that compensatory regulation also occurs in the opposite direction.

At gene promoters, H3K4me3 depletion in H3K4M EpiLC lines revealed distinct contributions of H3K4me3 and 5meC in restricting H3K27me3. H3K4me3 appears to protect the TSS from H3K27me3 accumulation at bivalent promoters, consistent with previous studies (Mas et al. 2018; Yagi et al. 2025). Targeted H3K4me3 demethylation at candidate bivalent promoters using the H3K4 demethylase KDM5B recapitulated the genome-wide observation that H3K27me3 accumulates upon loss of the permissive mark. Relatively few DMRs were called at promoters, and fewer still were associated with gene misregulation (Fig. S8D). However, 5meC presence across gene bodies and at bivalent promoter shores could limit the spreading of H3K27me3 (Fig. 7K, bottom) (Wu et al. 2010; King et al. 2016; Richard Albert et al. 2025). Curiously, we noted a stronger H3K27me3 signal at bivalent CGI promoters in WT H3K4M EpiLCs compared to TKO H3K4M (Fig. 6C), despite a greater reduced promoter accessibility (Fig. 6F; Fig S8G,H) and increased number of downregulated genes (Fig. 4E,F) in the latter. This would argue against increased H3K27me3 enrichment at TSSs, *per se*, being the most significant determinant of gene repression. One possibility is that in the TKO H3K4M EpiLCs, the spreading of PRC2 into gene bodies creates features akin to “Polycomb domains” that are less permissive to transcription factor binding and gene expression (Blackledge and Klose 2021).

With these two models established, we can posit whether enhancer or promoter misregulation is the more important factor in the global disruption of gene expression in the TKO H3K4M line. There is compelling evidence that enhancers are the primary underlying driver. Firstly, H3K4M induction has a substantially stronger effect on H4K4me1 levels compared to H3K4me3. Given that H3K4me1 is the enhancer-associated mark, this would indicate that enhancers are more sensitive to H3K4M transgene expression. Consistently, in naïve ESCs, we detected a particularly small effect on H3K4me3 levels at promoters, which was discordant with the far greater number of differentially expressed genes. Secondly, a direct effect of 5meC is more plausible at enhancers, where DNA methylation levels are generally intermediate, than at most promoters, where 5meC is essentially absent. Moreover, H3K4me1 and 5meC can coexist—at least to an extent (Morales-Franco et al. 2026); in contrast, H3K4me3 and 5meC are mutually exclusive (Sharifi-Zarchi et al. 2017; Otani et al. 2009). Therefore, the presence of both H3K4 and DNA methylation at enhancers can more reasonably cooperate to protect from Polycomb nucleation. This was borne out in the data, where we observed a pronounced gain of H3K27me3 marks in the TKO H3K4M EpiLCs. Finally, chromatin changes at promoters could be a downstream consequence of repressed enhancer activity. However, it is important to note that the two models are not necessarily incompatible. Furthermore, the relationship between chromatin state and transcription is likely to be context-dependent, with alterations in accessibility, histone modifications, enhancer activity, and transcription potentially reinforcing one another via different means depending on the locus examined.

A striking finding from this study is the unexpected cellular phenotypes exhibited in the TKO H3K4M line. Notably, there was an absence of overt phenotypic defects in naïve ESCs. It is likely not a coincidence that even in WT conditions, global DNA methylation levels are low in the naïve state. Therefore, naïve ESCs expressing H3K4M can tolerate or buffer transcriptional dysregulation while maintaining pluripotency and self-renewal. However, in contexts where 5meC levels are globally high—serum ESCs, EpiLCs and EBs—the reduction or absence of both 5meC and H3K4me was catastrophic. The basis of this observation remains unclear. Depending on the cellular context examined, GSEA revealed terms that may provide insight into the underlying causes of the observed phenotypes, e.g., developmental pathways, immune response genes, and apoptosis. However, the GSEA terms were not uniform across experiments. Furthermore, it is difficult to disentangle the cause from consequence when genes are differentially expressed. For example, the apoptosis genes enriched in TKO H3K4M EpiLCs might be a response to other cellular abnormalities. It is also important to note that while this study focused on protein-coding gene regulation, the phenotypes could be related to compromised genome integrity or reduced control of the repetitive genome. Future work will be necessary to better understand the processes underlying the cellular phenotypes.

Finally, it is worth noting that the H3K4M transgene has been shown to destabilize MLL3/4 (KMT2C/D), the major depositors of H3K4me1 at enhancers (Jang et al. 2017, 2019). Such disruption could contribute to the defects observed in our system, as MLL3/4 regulate enhancer activity beyond their catalytic activity, notably through the recruitment of UTX (KDM6A), an H3K27 demethylase, and the histone acetyltransferase p300, which deposits H3K27ac (Wang et al. 2016; Dorighi et al. 2017). More broadly, although no comparable effects have been demonstrated for other H3K4 methyltransferases that deposit H3K4me3, we cannot rule out the possibility that H3K4M expression alters the recruitment, activity, or chromatin binding of these enzymes, with loss of H3K4 methylation being a consequence of their altered function rather than the sole driver of the observed phenotypes. Increasing evidence indicates that H3K4 methyltransferases can regulate transcription and chromatin function independently of their catalytic activity, raising the possibility that such functions may contribute to the phenotypes observed in our system (Au et al. 2025; Hoshii 2025). Nevertheless, our epigenome editing experiments using the KDM5B catalytic domain provide direct evidence that H3K4me3 itself contributes to the local regulation of H3K27me3 levels.

Overall, this work challenges our classical view of H3K4me and 5meC as strictly mutually exclusive. Instead, they appear to act cooperatively to maintain a permissive chromatin state and prevent excessive PRC2 repression. Together, these findings provide an integrated understanding of how epigenetic mechanisms collectively preserve transcriptional programs and cellular identity.

## Material and Methods

### Mouse embryonic stem cell culture

All mouse embryonic stem cell (ESC) lines in this study were derived from the E14Tg2a (E14) parental line. The *Dnmt* triple-knockout (TKO) line was previously generated in-house (Dubois et al. 2022). All ESCs were grown at 37°C and 5% CO_2_.

For serum culture conditions, cells were grown in ESC serum media, composed of Glasgow medium (Gibco), 15% fetal bovine serum (FBS), 0.1 nM MEM non-essential amino acids, 1nM sodium pyruvate, 2 mM L-Glutamine, penicillin/streptomycin, 0.1 nM β-mercaptoethanol and 1000 U/mL leukemia inhibitory factor (LIF). To pass the cells, medium was removed and a 1X PBS wash was performed before a 5 min incubation with Trypsin-EDTA 0.05% (Gibco) at 37°C. Trypsin was then neutralized with an equal volume of serum media and cells were resuspended to detach and disaggregate.

For 2i+VitC culture conditions, cells were grown in N2B27 medium (50% neurobasal medium and 50% DMEM) supplemented with N2 (Gibco), B27 (Gibco), 2 mM L-glutamine, 0.1 mM β-mercaptoethanol, penicillin–streptomycin, 1000 U/mL LIF, 2i (3 μM GSK3β inhibitor CT-99021 and 1 μM ERK inhibitor PD0325901) and Vitamin C (Sigma) at a final concentration of 100 μg/mL. To pass the cells, the medium was removed, Accutase (Gibco) was added and cells were incubated 5 min at room temperature to detach and disaggregate. Wash media (50% neurobasal medium and 50% DMEM) was used at a similar volume to detach and resuspend the cells. Cells were then centrifuged at 1200 rpm for 3 min, supernatant was removed and cells were resuspended in fresh Wash media before being seeded at the required cell concentration.

Cells cultured in serum or 2i+VitC media were grown on feeder-free 0.1% gelatin-coated flasks or plates.

For EpiLC differentiation, 2i+VitC cells were splitted, washed with Wash media and seeded at a density of 2×10^5^ cells per cm^2^ on fibronectin (10 µg/mL, Sigma)-coated plates or flasks and grown in N2B27 medium supplemented with 20 ng/mL Activin A (R&D) and 12 ng/mL FGF2 (R&D). To count or collect EpiLCs, cells were washed with Wash media and detached with Accutase in the same way as for the 2i+VitC cells.

When needed, cells were resuspended and counted using a Countess II cell counter (Invitrogen), with a 20 μL 1:1 mix of cell suspension and trypan blue. Then, the desired number of cells was seeded on a new plate or flask or was collected for experiments.

To perform doxycycline (Dox) induction in pTREG3 lines, cell culture media was supplemented with 0.5µg/mL of doxycycline, at least 24 h before the start of experiment. For degradation of the RbBP5-FKBP protein, dTAG-13 (Tocris Bioscience) was resuspended in DMSO and added at a final concentration of 100 nM in culture media at the start of the experiment.

GSK-3484862 (MedChemEXpress) at a final concentration of 2 µM was used for a minimum of four days to inhibit DNMT1 and reduce 5meC levels in WT pTREG3 cells.

To inhibit EZH2 and reduce H3K27me3 levels in pTREG3 lines, UNC 1999 (inhibitor; Tocris) and UNC 2400 (Mock; Tocris) were added in the cell culture media at a final concentration of 1 µM, for a minimum of seven days.

Mycoplasma contamination checks were performed using an ultrasensitive qPCR assay (Eurofins).

### Plasmid Generation

The inducible pTRE3G-H3.3-P2A-Ruby (H3) and pTRE3G-H3K4M-P2A-Ruby (H3K4M) were generated by modifying pAU003 (www.addgene.org/156430/). Briefly, pAU003 was digested by ClaI/AflII, and the H3 or H3K4M genes were inserted by Gibson cloning.

The constitutively expressed dCas9-SunTag-P2A-scFv-GFP-TET1CD (dCas9-ST/TET1) construct used for targeted epigenome editing was a gift from T. Horii and I. Hatada (Horii et al. 2022). This system was adapted to generate dCas9-ST/KDM5B wild-type (WT) and catalytically inactive (CI) constructs. KDM5B constructs were kindly given by M. S. Oak and E. Hörmanseder (Oak et al. 2025), and the WT or CI catalytic domain was amplified and introduced into the digested dCas9-ST vector by Gibson cloning, after removing the TET1 CD by AgeI/NotI digestion.

The multi-guide plasmid containing four sgRNAs targeting the locus of interest was generated as previously described, using adapted Golden gate assembly protocol described in (Genolet et al. 2021). Amplified and ligated sgRNA sequences were assembled in a piggyBac-compatible vector that contains both Ruby fluorescence and a Hygromycin resistance marker. All guide sequences can be found in *Supplemental Table 1*.

Whole plasmid sequencing (Plasmidsaurus) was performed for all constructs used in this study.

### Transgenic cell line generation

All transfections were performed in serum+LIF-cultured ESCs (see Cell culture section) using Lipofectamine 2000 (Life Technologies) according to the manufacturer’s instructions.

The inducible pTREG H3 or H3K4M constructs were targeted to the *Tigre* safe-harbour locus in combination with a modified pX330 (www.addgene.org/42230) in order to generate a CRISPR/Cas9-mediated double-stranded break. Following transfection, cells were selected with 1 µg/mL puromycin for 2 days starting 24 h post-transfection. Individual colonies were isolated, expanded, and validated by genotyping. Three selected clones were further validated by western blot analysis. Unless otherwise indicated, one representative clone of each pTRE3G cell line was used for subsequent experiments.

The RbBP5-FKBP degron donor construct and corresponding CRISPR plasmid were kindly provided by H. Y. A. Au and R. J. Klose, and cell lines were established as previously described (Au et al. 2025). Independent RbBP5-FKBP degron clones were used as biological replicates.

To generate epigenome editing cell lines, WT H3K4M cells were co-transfected with the multiple-guide-containing plasmid, the dCas9-ST/TET1 construct, and the piggyBac transposase plasmid pBroad3_HyPBase_IRES_RFP. Scrambled control cells were transfected without the multiple-gRNA plasmid, as scrambled gRNAs are present within the epigenome editing construct itself, as previously described (Richard Albert et al. 2025). Using the same strategy, WT H3 and TKO H3 cells were co-transfected with either the dCas9-ST/KDM5B WT or CI constructs.

Twenty-four hours post-transfection, cells that received the multiple-guide RNA plasmid were subjected to hygromycin selection (200 µg/mL; Invitrogen) for 11 days. Scrambled control cells were not hygromycin-selected. Four to six days after transfection, GFP-positive cells were sorted for scrambled control conditions, and GFP/Ruby double-positive cells were sorted for targeted conditions, as the multi-guide plasmid co-expresses the Ruby fluorescent reporter. Cell sorting was performed using a BD FACSAria™ Fusion cell sorter (Becton Dickinson). Sorted cells were expanded and used for subsequent experiments.

All sgRNA sequences were designed using the CRISPOR web tool (Concordet and Haeussler 2018) and are listed in *Supplemental Table 1*.

### ESC growth assay

To perform cell growth assays in serum and 2i+VitC culture conditions, 7.5×10^5^ cells per well were plated in a 6-well plate. Media was changed every day until the end of the experiment on the sixth day. Every two days, cells were detached, resuspended and counted using the Countess II cell counter (Invitrogen) as described above. Cells were seeded again at 7.5×10^5^ cells per well. Cell growth was normalized by multiplying the measured number of cells by the ratio of the previous cell count to the reseeded cell number, resulting in a cumulative growth curve.

EpiLC growth assay was started by seeding 2×10^5^ cells per well in a 6-well plate for each time point on day 0. Cell growth was assessed every 24 h until the end of the experiment on day 4.

### Alkaline-phosphatase colony formation assay

Before starting the experiment, 6-well plates were coated overnight with poly-L-ornithine 0.01% at 37°C. The next day, the poly-L-ornithine was removed and washed twice with 1X PBS, and then wells were coated with 1X laminin for 2 h at 37°C. Before seeding them, cells were trypsinized and counted, and adequate serial dilutions were performed to seed 300 cells per well. Media (serum+LIF or serum-LIF) was changed once or twice during the duration of experiment when needed. On the seventh day, cells were washed once with 1X PBS, and fixed and stained using the leukocyte alkaline phosphatase kit (Sigma) or using NBT/BCIP diluted stock solution (Roche). For Sigma’s kit, the citrate-acetone-formaldehyde and staining solutions were prepared according to the manufacturer instructions. 1 mL of the fixing solution was added per well for 1 min to fix the cells, then removed. Cells were washed with H^2^O MilliQ. Finally, 1 mL of the staining solution was added per well. After 20 min in the dark, cells were washed again with H_2_O MilliQ and left to dry at room temperature. For NBT/BCIP staining procedure, 10 mL of staining buffer was freshly prepared (0.1 M Tris-HCl pH 9.5, 0.1 M NaCl, 0.05 M MgCl_2_ and H_2_O MilliQ), supplemented with 200 µL of NBT/BCIP stock solution. The solution was kept in the dark. After fixation, cells were washed three times with 1X PBS, and 1 mL of staining solution was added per well for 15 min in the dark. Then, fixed cells were washed with 1X PBS and left to dry at room temperature.

Colonies were counted on multiple occasions by different experimenters whenever possible. Colony pluripotency status was assessed based on colony staining patterns, with pink (Sigma) or purple (NBT/BCIP) colonies classified as undifferentiated, colonies displaying staining restricted to the center with unstained edges classified as mixed, and unstained colonies classified as differentiated.

### Embryoid body differentiation and area measurement

Embryoid body (EB) differentiation was started from serum+LIF cultured cells. The day before starting the differentiation, 4×10^6^ cells were plated in ESC serum media in a 10 cm gelatinized plate. For the rest of the experiment, EB medium was used, composed of Glasgow medium (Gibco), 10.25%, FBS, 0.1 nM MEM non-essential amino acids, 1 nM sodium pyruvate, 2 mM L-Glutamine, penicillin/streptomycin and 0.1 nM β-mercaptoethanol. On day 0, 1×10^6^ cells were counted and resuspended in 20 mL of media. Using a multichannel pipette, rows of 20 µL drops containing approximately 1000 ESCs were aligned on the lid of a 15 mL plate containing 15 mL 1X PBS at the bottom. The dish was carefully closed and placed back in the incubator. The droplets remained hanging on the inside of the lid of the plate for the cells to aggregate. On day 2, EBs were collected, resuspended and transferred to a new non-adherent 10 cm non-treated plastic dish with fresh media. Every two days, EBs were transferred to a new dish with fresh media until the end of the experiment on day 8.

During differentiation, two replicates per condition were taken for area measurement every two days, starting at day 2 of differentiation. Pictures were taken using a EVOS XL Core inverted microscope (4X) (ThermoFisher). The EB area measurement was performed using FIJI. At least 5 independent EBs per replicate were measured at each timepoint. After measurements, the EBs were stored for subsequent RNA extraction and analyses (below).

### S2 drosophila cell culture

S2 drosophila cells were kindly provided by D. Holoch (Institut Curie) and were used as spike-in material for normalization of H3K27me3 CUT&Tag datasets. S2 cells were cultured at room temperature (∼25°C) in Schneider’s Drosophila medium supplemented with 10% fetal bovine serum (FBS) and penicillin/streptomycin. Cells were seeded at a density of 1×10^6^ cells/mL and passaged when reaching a density of 6–20×10^6^ cells/mL. The culture medium was replaced every 2–3 days. For cell collection, cells were counted, and the required number of cells was collected by centrifugation at 1000 rpm for 3 min to remove the culture medium.

### Protein extraction and immunoblot

Cell pellets were resuspended in a BC250 lysis solution (25 mM Tris pH 7.9, 0.2 mM EDTA, 20% glycerol and 0.25 M KCl) supplemented with protease inhibitors (Roche), and sonicated three times for 5 s each using a Bioruptor (Diagenode) on the high settings. This was followed by a 10 min centrifugation at 10,000 rpm and 4°C, then supernatant was transferred to a new tube. Pierce BCA protein assay kit (Thermo Fisher Scientific) was used for quantification of protein concentrations, with an Infinite M200 plate reader (Tecan). Protein samples were then prepared with 4X Laemmli loading buffer (Bio-Rad) and denatured at 95°C for 5 min. Samples were run at 150 V for 1 to 2 h on homemade Tris-Glycine SDS-PAGE gels containing 8% to 15% acrylamide, depending on the molecular weight of the protein of interest, preceded by a 4% stacking gel. After the migration, proteins were transferred onto a PVDF membrane using a semi-dry transfer system (Transblot Turbo, Bio-Rad). Ponceau staining was performed to assess protein transfer efficiency. H_2_O MilliQ was used to reveal staining, and remaining Ponceau was removed by two washes of PBS-Tween (1%) (PBS-T). Blocking was performed for at least 30 min in 5% milk in PBS-T. Next, membranes were incubated in primary antibodies diluted in 5% milk at 4°C overnight. A list of primary antibodies and associated dilutions is provided in *Supplemental Table 2*. The next day, membranes were washed three times with PBS-T for 15 min each, then incubated for 1 to 2 h in secondary antibody diluted in PBS-T. For all primary antibodies except anti-H3K27me3, the membranes were incubated in the dark with a fluorescent-tagged secondary antibody (Goat anti-Rabbit IgG StarBright^TM^ Blue 700, 1:2500). For anti-H3K27me3 detection, membranes were incubated with an HRP-conjugated secondary antibody and revealed using an enhanced chemiluminescence (ECL) substrate (Cytiva). Membranes were washed three additional times, and blots were imaged using a ChemiDoc system (Bio-Rad).

### Mass Spectrometry

To quantify histone post-translational modifications in pTRE3G cell lines, cells were cultured in serum+LIF conditions and collected after 24 h of Dox induction. Cells were washed three times with ice-cold 1X PBS and centrifuged at 1300 rpm for 3 min at 4°C. Cell pellets were snap-frozen in liquid nitrogen and stored at −80°C until shipment to the Protein Analytics Unit of the Biomedical Center, Ludwig Maximilian University of Munich. Sample preparation, LC-MS/MS acquisition, and primary data analysis were performed by the facility according to their established protocols (Völker-Albert et al. 2018; Alabert et al. 2020; Schäfer et al. 2026).

Briefly, pellets were resuspended in 0.2 M H_2_SO_4_ and incubated overnight at 1000 rpm and 4°C to perform complete histone extraction, as previously described (Schäfer et al. 2026). Following extraction, the samples were centrifuged at 15,000 rcf for 30 min at 4°C. Prior to tryptic digestion, lysine residues were propionylated by incubating samples with 34.2 mg/mL NHS-propionate each in ACN for 30 min at 50°C. A fixed amount of stable isotope labelled histone peptides (modified and unmodified, label: R (15N, 13C); 500 fmol/µL, JPT Peptide Technologies GmbH, Germany) was added to the reaction mix, prior to digestion as quality control. Samples were digested with trypsin (0.5 ng/µL, sequencing grade, Promega) overnight at 37°C and 1000 rpm. Tryptic peptides were desalted and purified using C8 Stagetips. Samples were lyophilized in a SpeedVac until dryness. For analysis, samples were resuspended in 15 μL of 2% ACN, 0.3% TFA (PreOmics GmbH LC Loading buffer) and stored at -20°C until MS analysis or directly subjected for LC-MS/MS measurement.

1 μL was injected in an Ultimate 3000 RSLCnano system (ThermoFisher Scientific) and separated in a 25 cm analytical column (75 µm ID, 1.6 µm C18, Aurora-IonOpticks) with a 90 min gradient from 3 to 43% ACN in 0.1% formic acid at a flowrate of 300 nl/min before being electrosprayed into QExactive HF mass spectrometer (ThermoFisher Scientific). Survey full-scan MS spectra (from m/z 250 to 900) were acquired with resolution R = 60,000 at m/z 400 (AGC target of 3e6). Top 10 most intense ions (charge state between 2 and 5) were sequentially isolated with an isolation window of 2.0 m/z to a target value of 2e5 and fragmented at 27% normalized collision energy and acquired with resolution R = 15,000 at m/z 400 (max IT 60 ms). Ion spray voltage was set to +1.5 kV with no sheath and auxiliary gas flow and heated capillary temperature was set to 250°C.

Raw data from mass spectrometry was analyzed using the Skyline software (MacCoss Lab, University of Washington, Seattle, WA, USA) (Pino et al. 2020). Automatic selection of doubly and triply charged peptide masses was manually curated based on the relative retention times and fragmentation spectra. Integrated peak areas were exported and used to calculate relative levels of each PTMs using R environment (R Core Team, 2022), based on the formula given in (Völker-Albert et al. 2018).

### LUminometric Methylation Assay (LUMA)

Genomic DNA was extracted from collected cell pellets using the NucleoSpin Tissue kit (Macherey-Nagel). 500 ng of genomic DNA was digested with either MspI + EcoRI or HpaII + EcoRI (New England BioLabs), with EcoRI included as an internal normalization. The enzymatic digestions were quantified by pyrosequencing using a PyroMark Q24 system and global CpG methylation levels were calculated based on the normalized peak height ratio of HpaII to MspI.

### Bisulfite-pyrosequencing (BS-pyro)

Bisulfite conversion was performed on 500 ng of genomic DNA using the EZ DNA Methylation-Gold kit (Zymo). PCR was performed on target genomic regions with Immolase (Ozyme) polymerase and with primer pairs recognizing BS-converted DNA, including a biotin-conjugated primer, starting from 1 µL of bisulfite-converted DNA. Primer pairs are found in *Supplemental Table 1*. The PyroMark Q24 Advanced reagents (Qiagen, 970922) were used to prepare sequencing reactions according to the manufacturer’s instructions. Sequencing was performed by the PyroMark Q24 pyrosequencer (Qiagen) and the PyroMark Q24 Advanced 3.0.1 software was used to design the Pyrosequencing assays and analyze the results.

### ChIP-qPCR

ChIP-qPCR was performed as previously described, with minor modifications (Monteagudo-Sánchez et al. 2024). Briefly, 5 to 9×10^6^ cells were collected, crosslinked using formaldehyde (1%) and lysed in lysis buffer (10 mM TRIS pH 8, 100 mM NaCl, 1 mM EDTA pH 8, 0.5 mM EGTA pH 8, 0.10% Sodium Deoxycholate, 0.5% N-lauroylsarcosine and protein inhibitors) for 10 min at 4°C on a rotator. Chromatin was sonicated four times during 10 min on high settings with 30 sec ON/30 sec OFF program, and checked on agarose gel for 200-700 bp enriched fragments. In parallel, 50 µL of protein A magnetic beads (Invitrogen 10001D) were incubated with primary antibody. List of antibodies and concentrations are found in *Supplemental Table 2*. Following incubation, beads coupled to antibodies were added to the sonicated chromatin and incubated overnight at 4°C. The next day, beads were washed twice with TF-WBI buffer (20 mM Tris–HCl/pH 7.4, 150 mM NaCl, 0.1% SDS, 1% Triton X − 100, 2 mM EDTA), twice with TF-WBIII buffer (250 mM LiCl, 1% Triton X-100, 0.7% DOC and 10 mM Tris–HCl, 1 mM EDTA), and twice with TET buffer (0.2% Tween-20, 10 mM Tris–HCl/pH 8.0, 1 mM EDTA). Next, chromatin elution and de-crosslinking was performed during overnight incubation at 65°C in 70 µL of elution buffer (0.5% SDS, 300 mM NaCl, 5 mM EDTA, 10 mM Tris–HCl pH 8.0) containing 40 µg of proteinase K. DNA was eluted and purified using the MinElute Qiagen kit (28206), and used for qPCR. Enrichment was calculated relative to input (1%). Primers sequences can be found in *Supplemental Table 1*.

### RNA extraction

Total RNA extraction from cell pellets was performed using the KingFisher Duo Prime Magnetic Particle Processor associated with the MagMAX mirVana Total RNA kit, or using the NucleoSpin RNA kit (Macherey-Nagel) according to the manufacturer’s instructions.

### RT-qPCR

From extracted RNAs, first strand cDNA synthesis was performed with the SuperScript III Reverse Transcriptase kit (Invitrogen). When possible, 500 ng of RNA were used, otherwise the highest volume of sample allowed (11 µL) was used for the reaction, along with 1 µL of 50 ng of random primers, 1 µL of 10 mM dNTP mix and nuclease-free H_2_O up to 13 µL. The rest of the procedure was done following the manufacturer’s instruction.

RT-qPCR reactions were composed of 0.5 µL of cDNA mixed with 5 µL of LightCycler 480 SYBR Green I Master, 0.5 µL of 10 µM of each forward and reverse primers (provided in *Supplemental Table 1*) and sterile H_2_O up to 10 µL.

The reaction was run on a LightCycler 480 II (Roche Applied Science) using a 384 well plate. The samples first followed an initial incubation at 95°C for 10 min, and then 45 cycles of denaturation at 95°C for 10 s, annealing at 61°C for 20 s and extension at 72°C for 20 s. Samples were amplified in triplicates with appropriate non-template controls. Relative gene expressions were calculated using the 2 −ΔΔC_t_ method and normalized to the geometric mean of the C_t_ value expression levels of the two housekeeping genes *Rrm2* and *Rplp0*.

### RNA-seq

Total RNA was sent to Novogene for poly(A)-selected RNA library preparation and sequencing on an Illumina NovaSeq X (150 bp paired-end reads; PE150).

### WGBS

Genomic DNA was extracted using phenol:chloroform precipitation. WGBS library preparation and paired-end sequencing (PE150) were performed by Novogene using the Illumina NovaSeq X Plus platform.

### CUT&Tag

Unless otherwise specified, steps were performed at room temperature.

CUT&Tag was performed as previously described (Morales-Franco et al. 2026; Kaya-Okur et al. 2019) with minor modifications. 5×10^5^ cells were collected. The CUT&Tag-IT® Spike-In Control (Active Motif) was used as an internal spike-in control for H3K4me1 and H3K4me3 CUT&Tag experiments, at a ratio of 1:20 (25,000 nuclei). After cell collection, nuclei were extracted from ESCs prior to mixing with spike-in control nuclei. Briefly, cells were incubated in 200 µL of ice-cold nuclei extraction (NE) buffer (20 mM HEPES-KOH pH 7.9, 10 mM KCl, 0.1% Triton X-100, 20% glycerol, 0.5 mM spermidine, and 1X protease inhibitor cocktail) on ice for 10 min. Nuclei were then centrifuged for 3 min at 600 × g and resuspended in 100 µL of ice-cold NE buffer before being mixed with spike-in control. For H3K27me3 CUT&Tag experiments, spike-in normalization was performed using S2 drosophila cells cultured in-house (see above). A total of 2×10^5^ S2 cells were mixed directly with ESCs prior to CUT&Tag processing.

Cells were washed once with 1 mL of Wash Buffer (20 mM HEPES-KOH pH 7.5, 150 mM NaCl, 0.5 mM Spermidine, 1X Protease inhibitor cocktail), and then resuspended in 200 μL of Wash Buffer. In parallel, 10 μL of Concanavalin A-coated magnetic beads (BioMag®Plus) were activated in Binding Buffer (20mM HEPES pH 7.5, 10 mM KCl, 1 mM CaCl_2_, 1 mM MnCl_2_), and mixed with the cells or nuclei on a rotator for 10 min. Tubes were then placed on a magnet and supernatant was discarded. Beads bound with cells were resuspended in 100 μL of Antibody Buffer (20 mM HEPES pH 7.5, 150 mM NaCl, 0.5 mM Spermidine, 1X Protease inhibitor cocktail, 0.01% Digitonin, 2 mM EDTA, 0.1% BSA) supplemented with primary antibody (*Supplemental Table 2*). Cells were incubated overnight at 4°C on a rotator. Then, tubes were placed on magnet to remove supernatant, and beads with cells were resuspended in 100 μL of Antibody Buffer containing Guinea Pig anti-Rabbit IgG antibody (ABIN101961) at 1:100 dilution, and incubated for 30 min on a rotator. Cells bound to the beads were washed three times with 1 mL Dig-Wash Buffer (20 mM HEPES pH 7.5, 150 mM NaCl, 0.5 mM Spermidine, 1X Protease inhibitor cocktail, 0.01% Digitonin) and incubated with pA-Tn5 loaded with methylated adapter (detailed in (Morales-Franco et al. 2026)) at 1:200 dilution in 100 μL Dig-300 Buffer (20 mM HEPES pH 7.5, 300 mM NaCl, 0.5 mM Spermidine, 1X Protease inhibitor cocktail, 0.01% Digitonin) for 1 h on a rotator. After, the bead-bound cells were washed three times with 1 mL Dig-300 Buffer in order to remove unbound pA-Tn5. Then, pA-Tn5 was activated for tagmentation by resuspending the cells on beads in 100 μL of Tagmentation Buffer (20 mM HEPES pH 7.5, 300 mM NaCl, 0.5 mM Spermidine, 1X Protease inhibitor cocktail, 0.01% Digitonin, 10 mM MgCl_2_), incubated at 37°C and at 1000 rpm for 1 h. DNA was purified with the Qiagen MinElute kit (28206) according to the manufacturer’s instructions. Oligonucleotide replacement and gap repair was performed as previously reported (Kaya-Okur et al. 2019; Lhoumaud et al. 2019). Briefly, 11 μL of the CUT&Tag DNA eluate was mixed with 2 μL of 10 μM Tn5mC-Repl01 oligo, 2 μL of 10X ampligase buffer, 2 μL of dNTPs at 2.5 mM and incubated in a PCR thermocycler with the following program: 50°C for 1 min, 45°C for 10 min, ramp down to 37°C at a rate of −0.1°C/s, hold at 37°C. Then 1 μL T4 DNA polymerase and 2.5 μL ampligase were added without removing the tube from the thermocycler and the reaction was incubated at 37°C for 30 min, then held at 4°C. Next, DNA was PCR-amplified and barcoded in 30 μL reactions (15 μL 2x KAPA HiFi HotStart Uracil+ ReadyMix, 13 μL eluted CUT&Tag DNA, 1 μL 10 μM i5 index primer, 1 μL 10 μM i7 index primer), with the following PCR thermocycler program: 72°C for 5 min; 98°C for 30 s; 12 cycles of 98°C for 10 s for M-ATAC or 14 cycles of 98°C for 10 s for M-CUT&Tag, 63°C for 30 s and 72°C for 30 s; and a final elongation at 72°C for 1 min. DNA was then purified using SPRI AMPure XP beads with a bead-to-sample ratio of 1.8:1 (no size selection) and eluted in 25 μL of H_2_O. An Agilent 2200 TapeStation with a D1000 screentape was used to determine concentration and size distribution of the pre-sequencing library. DNA libraries were sequenced paired-end (PE150) on a NovaSeq6000 instrument (Novogene).

### Methyl-ATAC-seq (M-ATAC)

Methyl-ATAC-seq libraries were generated, but only the ATAC-associated sequencing data were analyzed in this study.

M-ATAC was performed as previously described (Morales-Franco et al. 2026; Lhoumaud and Skok 2021; Lhoumaud et al. 2019). Briefly, 5×10^4^ cells were collected, washed in cold 1X PBS and resuspended in 50 µL of cold lysis buffer (10 mM Tris-HCl, pH 7.4, 10 mM NaCl, 3 mM MgCl_2_, 0.1% IGEPAL CA-630). Then, for tagmentation, lysate was mixed with 25 μL of newly prepared tagmentation buffer (10 mM Tris-HCl pH 8.0, 5 mM MgCl_2_ and 10% v/v dimethylformamide), 2.5 μL transposase containing the methylated adaptors (detailed in (Morales-Franco et al. 2026)), and 22.5 μL of nuclease-free H_2_O for 30 min at 37°C. DNA was purified with the Qiagen MinElute kit (28206), oligonucleotide replacement and gap repair was performed as described above (CUT&Tag section), and DNA was bisulfite-converted using the Zymo EZ DNA Methylation-Gold Kit (D5006) following manufacturer’s instructions. Then, DNA libraries were PCR-amplified, assessed using TapeStation and sequenced as for CUT&Tag samples (see above).

#### Bioinformatic analyses

##### RNA-seq processing

RNA-seq data was processed as previously described (Richard Albert et al. 2025). RNA-seq read quality was assessed using FastQC (version 0.11.9; (Andrews 2010)). PCR duplicate reads were removed using BBMap clumpify (version 38.18; (Bushnell 2014)) and the following parameters: reorder, dedupe = t, k = 19, passes = 6 and subs = 0.01 × read length, where the number of substitutions allowed is equal to the length of the read in nucleotides multiplied by the conservative sequencing error rate (1%) of Illumina machines. Lastly, adaptor and low-quality sequences were trimmed using Trimmomatic (version 0.39; (Bolger et al. 2014)) using the following parameters: ILLUMINACLIP:adaptors.fa:2:30:10 SLIDINGWINDOW:4:20 MINLEN:24, where adaptors.fa contains the FASTA sequence of Illumina sequencing adaptors. Reads were then aligned to the mm10 genome with STAR (version 2.7.9a; (Dobin et al. 2013)) using following parameters: --runMode alignReads --outFilterType BySJout -- outSAMtype SAM.

### CUT&Tag processing

PCR duplicates and low-quality reads were processed similarly to RNA-seq data (see above). Reads were aligned to the mm10 reference genome with Bowtie2 (version 2.4.5; (Langmead et al. 2019)) with parameters: --local --very-sensitive --no-mixed --dovetail --no-discordant -- phred33 -I 10 -X 700.

For H3K4me1 and H3K4me3 histone marks, BigWig files were normalized using exogenous spike-in reads mapped to the *D. melanogaster* genome (dm6). For H3K27me3 CUT&Tag datasets, spike-in normalization was performed using an approach adapted from Altamirano-Pacheco et al, in revision. The normalization factor (NF) was calculated as follows:

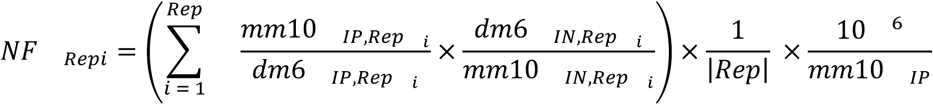

where Rep indicates the number of replicates per sample, mm10 refers to the mouse genome, and dm6 refers to the *Drosophila melanogaster* genome. IP represents the number of reads obtained from CUT&Tag sequencing, and IN represents the input normalization value estimated by qPCR amplification from genomic DNA.

Normalized BigWig files were generated with bamCoverage (version 3.5.1; (Ramírez et al. 2016)) function with the parameters: --scalefactor $SPIKE, --binSize 1, --smoothLength 0, --minMappingQuality 10, where $SPIKE corresponds to the normalization factor as defined above. BigWig files were merged by replicates with bigWigMerge (version 2; (Kent et al. 2010)) and converted with bedGraphToBigWig (version 4; (Kent et al. 2010)).

Peak calling for H3K4me1 and H3K4me3 histone marks was performed using macs2 (version 2.2.7.1; (Zhang et al. 2008)) callpeak with the following parameters: --broad, -f BAMPE, -g 1.87e9, -q 0.01, --nomodel and --shift 0. Blacklisted regions (defined by the Kundaje lab as part of the ENCODE consortium; (Amemiya et al. 2019)) as well as mitochondrial and non-canonical chromosomes were excluded from subsequent analyses. For H3K27me3, peak calling was performed with SEACR (version 1.3; (Meers et al. 2019)) with the parameters: 0.01, non, stringent. Peaks within a 10 kb interval were fused using BEDtools merge (version 2.31.1; (Quinlan and Hall 2010)) to generate H3K27me3 domains.

### M-ATAC processing

PCR duplicates were removed similarly to RNA-seq data (see above). Low-quality sequences were filtered with Trimmomatic using the following parameters: HEADCROP:18 ILLUMINACLIP:adapters.fa:2:30:10 SLIDINGWINDOW:4:20 MINLEN:24. Reads were then aligned to the mm10 mouse reference genome using Bismark (version 0.23.1; (Krueger and Andrews 2011)) with default parameters. Reads that lost their pair during filtering or that were not aligned in paired-end mode were re-aligned as single-end. Aligned reads were deduplicated with deduplicate_bismark.

Accessibility peaks were called with macs2 callpeak using the following parameters: -- broad, -f BAMPE, -g 1.87e9, -q 0.01, --nomodel and --shift 0. Blacklisted regions, mitochondrial and non-canonical chromosomes were excluded from subsequent analyses.

### WGBS processing

Low-quality reads were filtered using Trimmomatic and the following parameters: ILLUMINACLIP:adaptors.fa:2:30:10 SLIDINGWINDOW:4:20 MINLEN:36. Reads were aligned to the mm10 mouse reference genome using Bismark with default parameters. BigWig files of 5meC levels were generated from Bismark’s CpG report by extracting the genomic coordinates and methylation percentage of CpGs covered by at least five reads. The resulting file was transformed into bigWig using bedGraphToBigWig (version 4; (Kent et al. 2010)).

### Methylasso processing

Input files were obtained with the function coverage2cytosine from Bismark and the parameter:

--merge_CpG. Cytosines localised on non-canonical chromosomes were removed. Differentially methylated regions (DMRs) were identified with default MethyLasso (version 1.0.0; (Balaramane et al. 2024)) parameters.

### Profile plot and Heatmap generation

Promoter analyses were restricted to regions spanning ±1 kb around transcription start sites (TSSs) of UCSC refGene-derived transcripts.

CGI and non-CGI promoters were determined based on their overlap with known CpG islands, obtained from the “CpG Islands” track of the mm10 genome via UCSC Table Browser (Karolchik 2004). Overlaps were assessed with BEDTools intersect (version 2.31.1).

For profile plots spanning the TSS to TES (transcription end site) of genes, UCSC refGene transcript coordinates were obtained via the UCSC Table Browser.

Candidate enhancers for day 0 ESCs and day 2 EpiLCs were obtained from (Morales-Franco et al. 2026). Enhancers categorized as “othr” (no H3K4me1) in the corresponding day were removed from the analysis.

Signal quantification of histone marks for profile plots and heatmaps was performed with computeMatrix scale-regions (version 3.5.1) using the following parameters: -m 2000, -b 1000, -a 1000, --missingDataAsZero, --skipZeros, and filtering out blacklisted regions. DNA methylation quantification omitted the options “--missingDataAsZero, --skipZeros” and included the option “-bs 100”. Corresponding profiles were generated using plotProfile function (version 3.5.1). Heatmaps were generated with plotHeatmap (version 3.5.1).

### RNA-seq analysis

Expression values at the transcript (University of California, Santa Cruz (UCSC) RefGene) and gene (National Center for Biotechnology Information (NCBI) RefSeq) levels were calculated using VisR (version 0.9.42; (Younesy et al. 2015)) with uniquely aligned reads (MAPQ = 255). Alignment files were converted to BigWig using deepTools bamCoverage (version 3.5.1; (Ramírez et al. 2016)) and the following parameters: --binSize 1 --smoothLength 0 -- minMappingQuality 255 --normalizeUsing CPM --outFileFormat BigWig --blackListFileName ENCFF547MET.bed --ignoreForNormalization chrX chrM chrY. ENCFF547MET.bed corresponds to the coordinates of blacklisted regions defined by the Kundaje lab as part of the ENCODE consortium (Amemiya et al. 2019). Linear modeling using Limma (version 3.62.2; (Ritchie et al. 2015)) in R was used to identify statistically significant differential gene expression levels between samples. Voom normalization and a stringent threshold of |log2(fold-change)| > 1 and adjusted P-value (t-test) < 0.05 was applied to define statistical enrichment.

Gene Set Enrichment analysis (GSEA; (Mootha et al. 2003; Subramanian et al. 2005)) of mapped genes was performed with clusterProfiler (version 4.14.6; (Yu et al. 2012)) and visualized with enrichplot (version 1.26.6; (Yu 2026)). Gene ontologies depicted belong to "Biological Process".

UpSet plots were generated using the website: https://hiplot.cn/basic/upset-plot.

### Differential analyses of M-ATAC and CUT&Tag datasets

Differential analysis for chromatin accessibility over ATAC peaks was performed using R package DiffBind (version 3.20.0; (Stark and Brown)). Counting matrix was generated with the dba.count function using parameters: summits = FALSE, minOverlap = 2, bUseSummarizeOverlaps = FALSE, score = DBA_SCORE_READS_FOLD, filter = 5 and bParallel = FALSE. Data normalization was performed with dba.normalize using the DBA_DESEQ2 method and DBA_NORM_RLE normalization. Threshold for statistical significance (FDR) was 0.05. For H3K4me1 and H3K4me3, DiffBind was used similarly but without FDR threshold, and.bam files of reads aligned to the *D. melanogaster* genome (dm6) were also included to calculate spike-in normalization factors. The count matrix over histone modification peaks was calculated with dba.count with following parameters: summits = FALSE, bParallel=FALSE. Data normalization was performed using dba.normalize but with the argument spikein = TRUE.

For H3K27me3 differential analysis, a consensus domain set was generated by merging the H3K27me3 domains of all samples using BEDTools merge. The count matrix of the domain set was calculated with the function featureCounts from the R package Rsubread (version 2.24.0; (Liao et al. 2019)) with the following parameters: useMetaFeatures = FALSE, allowMultiOverlap = TRUE, isPairedEnd = TRUE, nthreads = 4. Data normalization was performed using the spike-in factor calculated above. Comparisons between conditions were performed with R package DESeq2 (version 1.50.2; (Love et al. 2014)).

For scatterplots, data from histone marks, accessibility and RNA-seq were associated to individual genomic regions with the function findOverlaps from R package GenomicRanges (version 1.58.0; (Lawrence et al. 2013)).

### Data visualisation

UCSC Genome Browser track hubs were used to visualize datasets at loci of interest.

Graphical representation and statistical analysis for growth curves, area measurements, colony formation assays, mass spectrometry, LUMA, pyrosequencing, and qPCR data were performed with GraphPad Prism (version 10.4.1).

Unless otherwise specified, bioinformatic analyses and data visualization were performed in R (version 4.4.2) and plotted with ggplot (version 4.0.3; (Wickham 2016)).

## Acknowledgements

We thank Nathalie Marcotte for experimental assistance and Priscillia Lhoumaud (IJM) for M-ATAC-seq and CUT&Tag expertise and advice. We thank Nicolas Valentin (IJM) from the flow cytometry platform at the Institut Jacques Monod for experimental assistance. We thank H. Y. A. Au and R. J. Klose (University of Oxford) for providing the RbBP5-FKBP degron and CRISPR constructs, T. Horii and I. Hatada (Gunma University, Japan) for providing the constitutive dCas9-SunTag/TET1CD epigenome-editing construct, O. Genolet and E. Schulz (MPI Berlin) for providing their protocol and plasmid templates for multiguide cloning, and M. Oak and E. Hörmanseder (Helmholtz Munich) for providing the KDM5B epigenome editing constructs. We thank D. Holoch (Institut Curie) for providing S2 drosophila cells. We thank A. Dubois and P. Navarro Gil (Institut Pasteur) for colony formation assay and embryoid bodies protocols, and E. Cazotes (Université Paris Cité) for NBT/BCIP staining protocol. We thank L. F. Altamirano Pacheco (Institut Pasteur) and D. Holoch (Institut Curie) for advice on H3K27me3 CUT&Tag spike-in normalization. This work was supported by the European Research Council (ERC-StG-2019 DyNAmecs) and by a Laboratoire d’Excellence Who Am I? (11-LABX-0071) emerging teams grant. A.D was supported by a 4th year PhD fellowship from the Fondation ARC; A.M-S. was supported by the Fondation pour la Recherche Médicale (FRM) Post doc France Fellowship (SPF202004011789) and Fondation ARC (ARCPDF12020070002563) postdoctoral fellowship; J.R.A was supported by the FRM Post doc France Fellowship (SPF202110014238).

## Author Contributions

Conceptualization: A.D., M.V.C.G.

Methodology: A.D., A.M-S., M.V.C.G.

Formal analysis: A.D., M.M-F., N.H., J.R.A., B.B.

Investigation: A.D., A.M-S., S.P., M.S., B.B., T.D.

Data curation: A.D., M.M-F., N.H., J.R.A.

Writing – original draft: A.D., M.V.C.G.

Writing – review & editing: A.D., M.M-F., M.V.C.G.

Supervision: M.V.C.G.

Project administration: M.V.C.G.

Funding acquisition: M.V.C.G.

## Data and code availability

Genomic data has been uploaded to the Gene Expression Omnibus (GEO), and is available upon request. Scripts for library processing can be found at https://github.com/julienrichardalbert/sra2bw. Scripts for data analysis and visualization can be found at https://github.com/Narjesse-hounani/K4M_project.

---

**Supplemental Figure 1.**
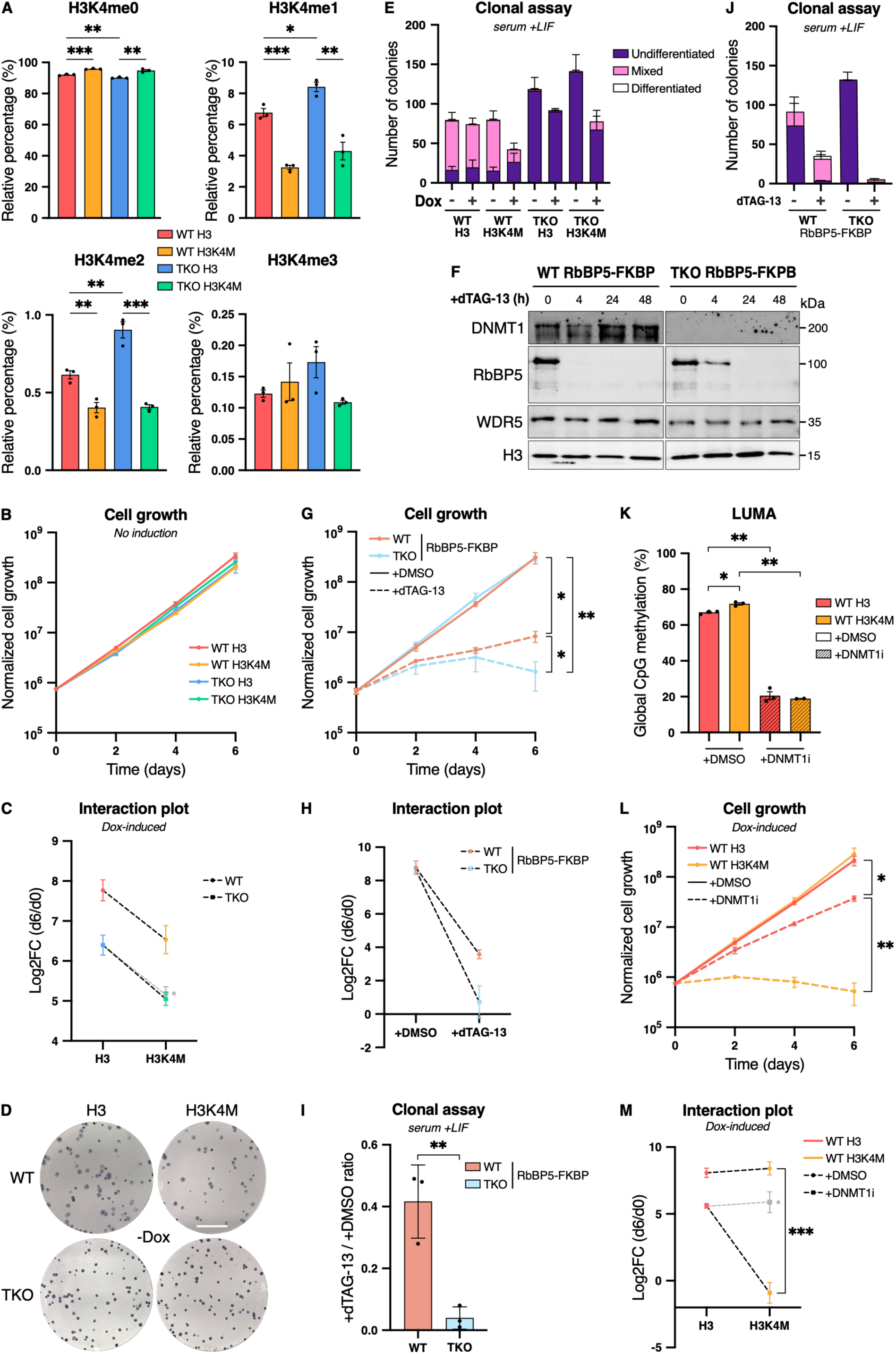
Loss of H3K4me and 5meC synergistically impairs proliferation in serum-grown ESCs. **A.** Mass spectrometry analysis of histone H3 peptides showing H3K4me fractions in all cell lines. Cells were grown in serum+LIF +Dox for 24 h. n = 3. B. Six-day growth assay in serum+LIF -Dox conditions. Normalized cell growth is represented in Iog10 scale, n = 3; except for WT H3, n = 2. C. Interaction plot representing the day 6 over day 0 ratio (Iog2) of cell numbers from serum+LIF +Dox six-day growth assay. * Grey line represents expected results if effects from 5meC and H3K4me depletion were additive, n = 3; except for WT H3, n = 5. D. Representative images of alkaline phosphatase-stained colony formation assays in serum+LIF -Dox conditions. The scale bar corresponds to 1 cm. **E.** Quantification of colony pluripotency types (undifferentiated, mixed, differentiated) in serum+LIF -/+Dox alkaline phosphatase-stained colony formation assays, n = 3. F. Immunoblot showing RbBP5 protein depletion in WT and TKO RbBP5-FKBP degron lines following dTAG-13 addition from 0 to 48 h, in serum+LIF culture conditions. G. Six-day growth assay for RbBP5-FKBP lines grown in serum+LIF conditions. +DMSO corresponds to the control without dTAG-13 addition. Normalized cell growth is represented in the Iog10 scale, n = 3; except for TKO +DMSO day 6, n = 2. H. Interaction plot representing the day 6 over day 0 ratio (Iog2) of cell numbers from serum+LIF six-day growth assay in RbBP5-FKBP degron lines. If RbBP5 depletion in TKO background led to additive effects only, the expected line would be as the WT line, n = 3; except for TKO +DMSO, n = 2. I. Ratio of alkaline phosphatase-stained colony counts in RbBP5-depleted (+dTAG-13) over control (+DMSO) conditions in serum+LIF. Replicates and their mean are shown, n = 3. J. Quantification of colony pluripotency types (undifferentiated, mixed, differentiated) in serum+LIF -/+dTAG-13 alkaline phosphatase-stained colony formation assays in RbBP5-FKBP lines, n = 3. **K.** LUminometric methylation assay (LUMA) representing global CpG methylation(%) in WT H3 and WT H3K4M cells grown in serum+LIF -Dox conditions and treated with DMSO (control) or DNMT1 inhibitor (+DNMT1Ì) for 4 days. Technical replicates and their mean are shown, n = 3; except for WT H3K4M +DNMT1Ì, n = 2. L. Six-day growth assay in serum+LIF +Dox conditions for WT H3 and WT H3K4M cells treated with DMSO (control) or DNMT1 inhibitor (+DNMT1Ì). Normalized cell growth is represented in the Iog10 scale, n = 3. **M.** Interaction plot representing the day 6 over day 0 ratio (Iog2) of cell numbers from serum+LIF +Dox six-day growth assay of WT H3 or H3K4M cells treated with +DMSO or +DNMT1Ì. * The grey line represents expected results if effects from 5meC and H3K4me depletion were additive, n = 3. **A-C, E, G-M:** Data are shown as mean ± s.e.m. of replicates from independent experiments. P-values were calculated using an unpaired Student’s t-test **(A, I),** on day 6 normalized cell counts **(G, L);** a paired Student’s t-test **(K)** or a two-way ANOVA test (M). * P≤0.05, ** P≤0.01, *** P≤0.001.

**Supplemental Figure 2.**
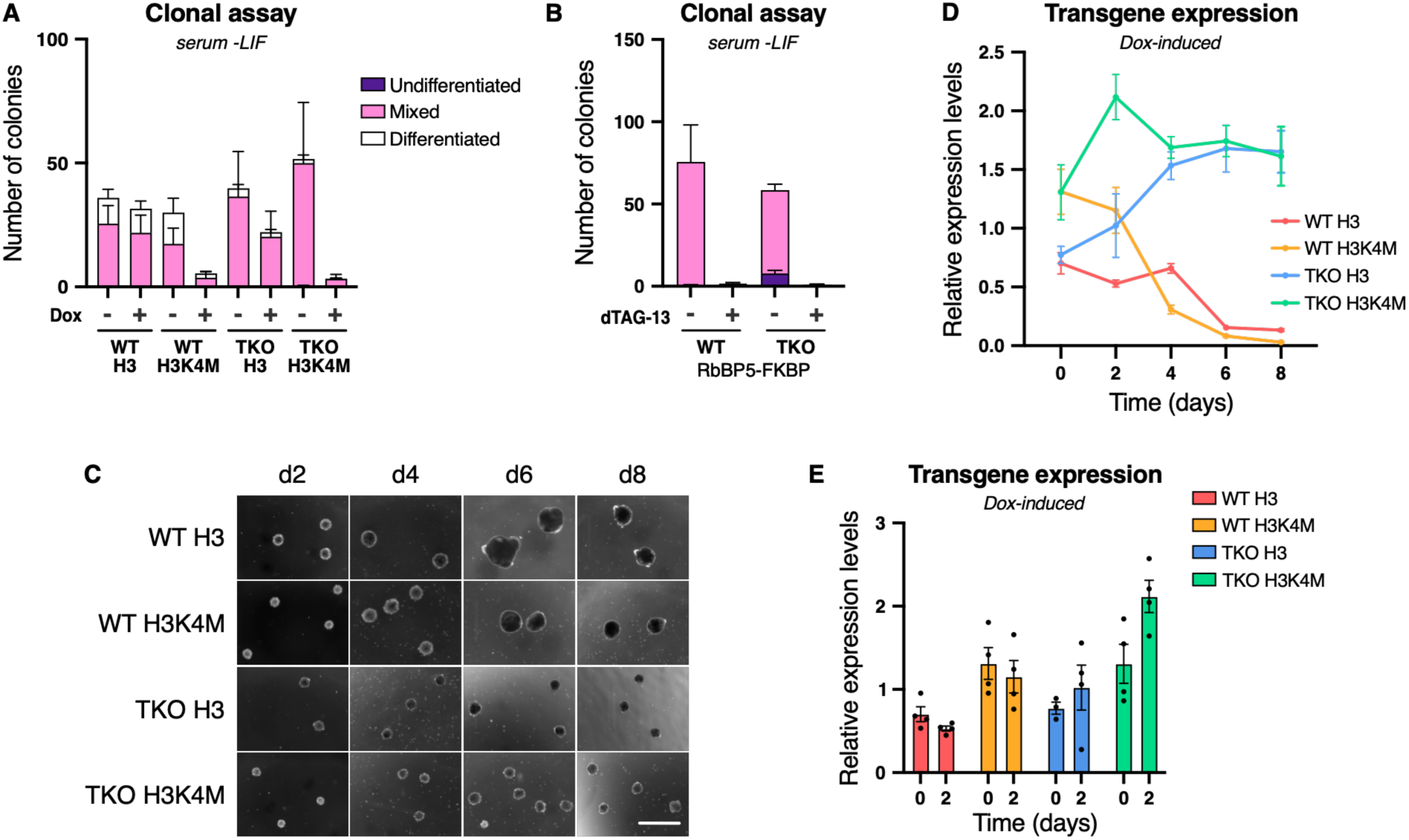
H3K4me is required for early-differentiation. **A.** Quantification of colony pluripotency types (undifferentiated, mixed, differentiated) in serum-LIF -/ +Dox alkaline phosphatase-stained colony formation assays in H3 and H3K4M lines, n = 3. **B.** Quantification of colony pluripotency types (undifferentiated, mixed, differentiated) in serum-LIF -/ +dTAG-13 alkaline phosphatase-stained colony formation assays in RbBP5-FKBP degron lines, n = 3. **C.** Representative images of embryoid bodies across differentiation from day 2 to day 8 for H3 and H3K4M lines (+Dox). The scale bar corresponds to 1 mm. **D, E.** Transgene expression from serum+LIF grown cells (day 0) to embryoid bodies until day 8 of differentiation (D) and from serum+LIF grown cells (day 0) and embryoid bodies day 2 (E), assessed by RT-qPCR. Relative expression levels are normalized to the average Ct of the two housekeeping genes *Rrm2* and *RpIpO* using the ΔCt method, n = 4 replicates obtained from two independent experiments. **A, B, D, E:** Data are shown as mean ± s.e.m. of replicates from independent experiments.

**Supplemental Figure 3.**
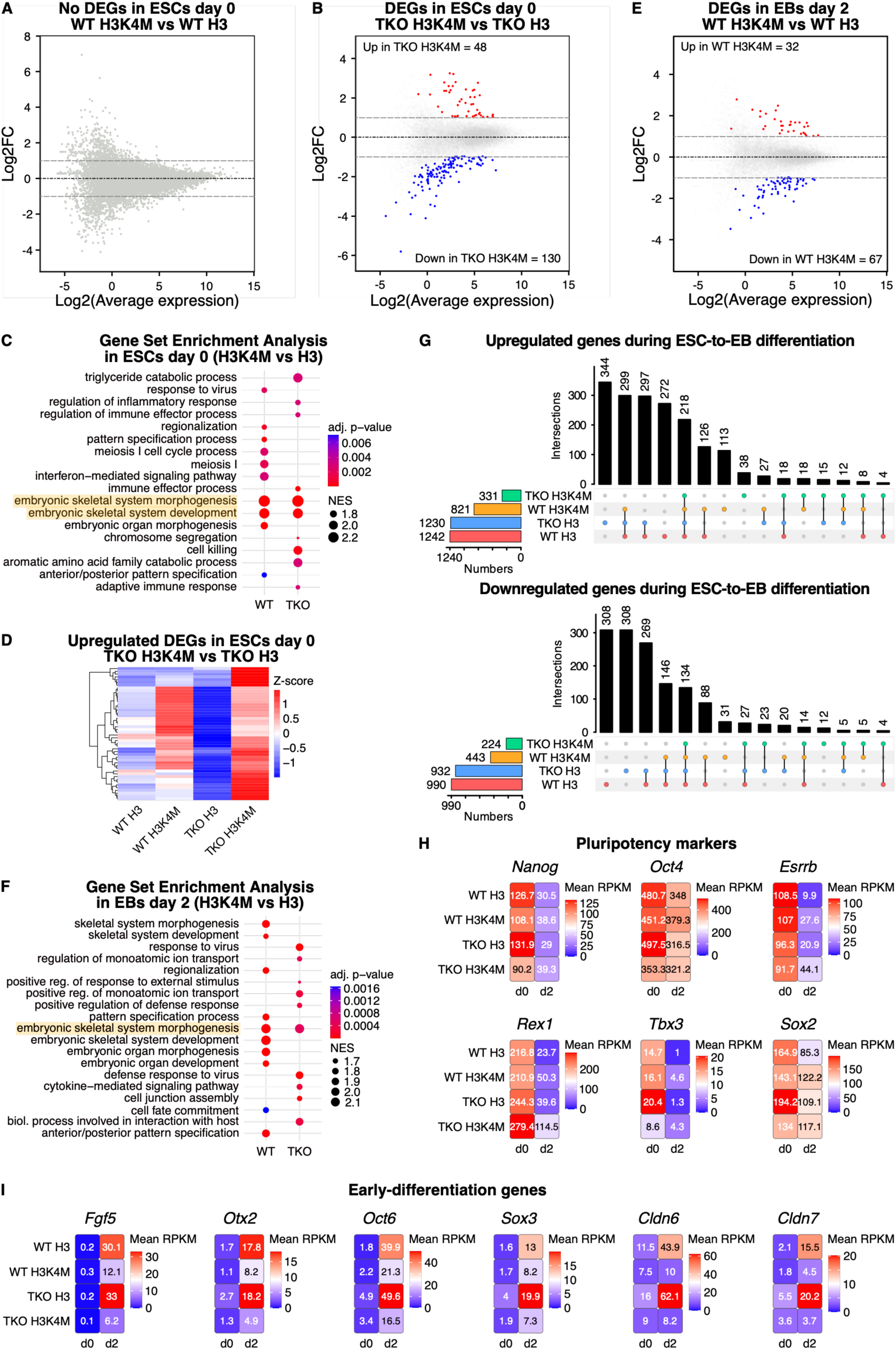
Loss of 5meC exacerbates H3K4M-induced transcriptomic defects in pluripotent cells and during embryoid body differentiation. **A, B.** MA plots representing differentially expressed genes (DEGs) (|log2FC| > 1 and adjusted p-value (t-test) < 0.05) in ESCs day 0 (serum+LIF +Dox) for WT H3K4M vs WT H3 (A) and TKO H3K4M vs TKO H3 (B) comparisons. The average expression is represented as Iog2. Numbers of significantly up- and downregulated genes are indicated **(B). C.** Gene Set Enrichment Analysis (GSEA) in ESCs day 0 (serum+LIF +Dox) for H3K4M vs H3 comparisons in WT and TKO genetic backgrounds. Highlighted terms are shared between both comparisons. NES = Normalized Enrichment Score. D. Heatmap of row-normalized gene expression (Z-score) for upregulated DEGs (log2FC > 1 and adjusted p-value (t-test) < 0.05; n = 48) in the TKO H3K4M vs TKO H3 comparison in day 0 samples. The mean of two replicates is shown. E. MA plot of differential gene expression (|log2FC| > 1 and adjusted p-value (t-test) < 0.05) in day 2 embryoid bodies (EBs) for WT H3K4M vs WT H3 comparison. The average expression is represented as Iog2. Numbers of significantly up- and downregulated genes are indicated. F. GSEA in day 2 embryoid bodies for H3K4M vs H3 comparisons in WT and TKO genetic backgrounds. The highlighted term is shared between both comparisons. G. UpSet plots representing the numbers of shared and unique upregulated (top) and downregulated (bottom) DEGs in embryoid bodies (day 2) vs ESCs (day 0) comparison for each cell line. **H, I.** Heatmaps showing mean expression levels (RPKM) of pluripotency markers (H) and early-differentiation markers (I) in ESCs (day 0) and embryoid bodies (day 2). The mean of two replicates is shown for each cell line and each time point.

**Supplemental Figure 4.**
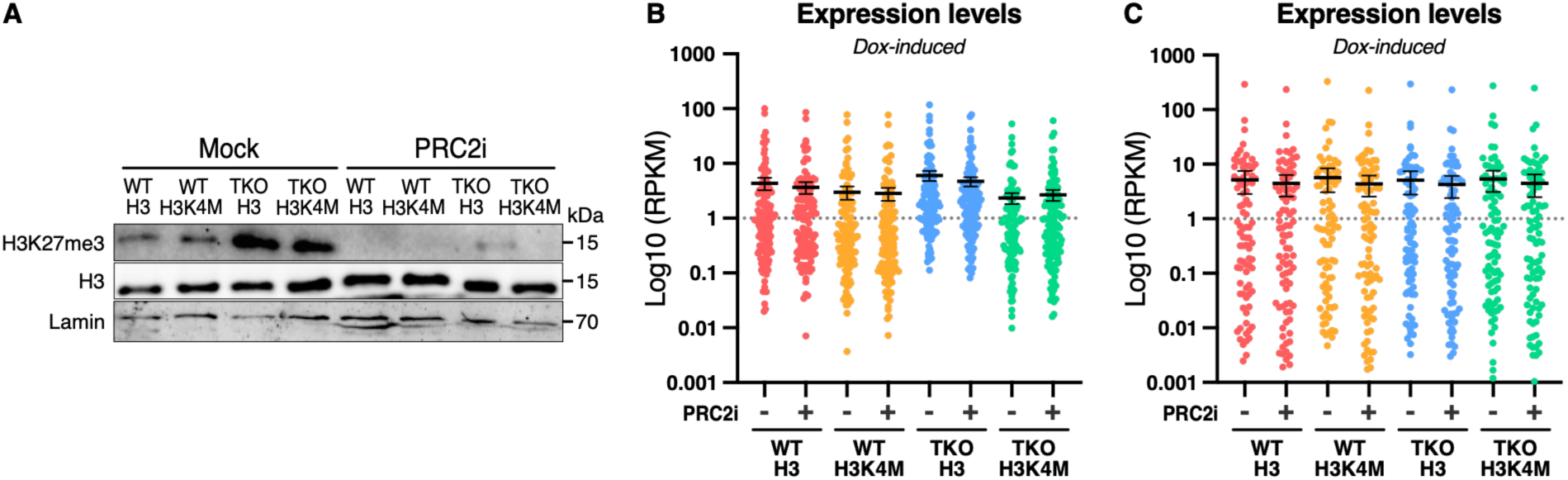
PRC2 inhibition reduces H3K27me3 levels and partially rescues downregulated gene expression in TKO H3K4M serum-grown cells. **A.** Immunoblot showing H3K27me3 levels in cells grown for 7 days in serum+LIF ÷Mock or +PRC2 inhibitor (PRC2Ì). B. Expression levels (loglO(RPKM)) of the 130 downregulated genes in TKO H3K4M vs TKO H3 day 0 comparison (no PRC2Ì; -PRC2Ì) in cell lines grown in -PRC2Ì or +PRC2Ì serum+LIF +Dox. **C.** Expression levels (loglO(RPKM)) of 130 randomly selected genes in cell lines grown in -PRC2Ì or +PRC2Ì serum+LIF +Dox. **B, C.** Each cell line in each condition is represented by three replicates. Data are shown as mean ± s.e.m..

**Supplemental Figure 5.**
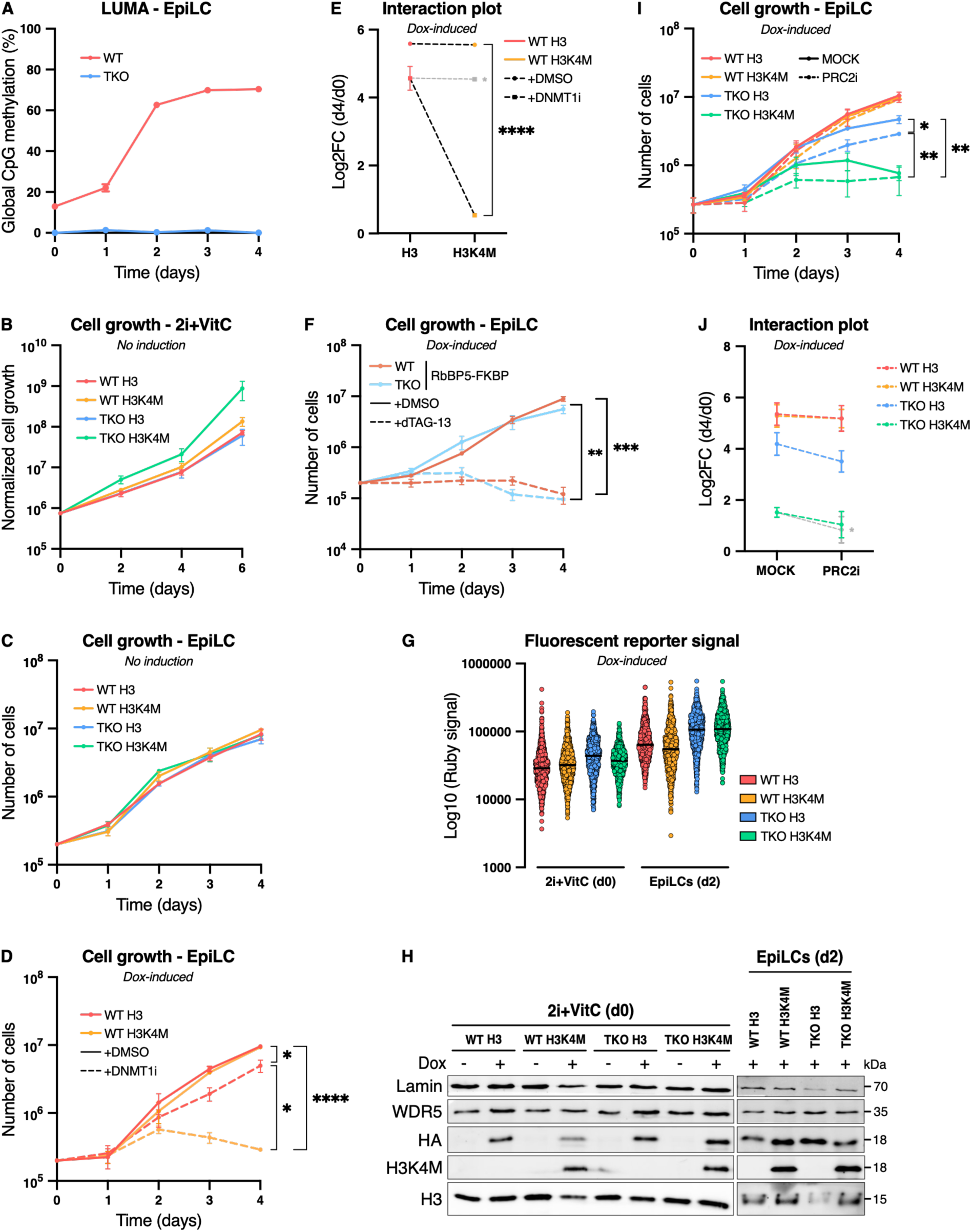
Loss of 5meC exacerbates EpiLC differentiation defects when H3K4me is depleted. **A.** LUMA representing global CpG methylation (%) in WT and TKO cells during four-day EpiLC differentiation (day 0 = 2i+VitC). n = 2 technical replicates; except for WT on day 3, n = 1. B. Six-day growth assay in 2i+VitC -Dox culture conditions. Normalized cell growth is represented in the Iog10 scale, n = 3. **C.** Four-day growth assay during EpiLC differentiation (-Dox). Cell counts are represented in the Iog10 scale, n = 3. D. Four-day growth assay during EpiLC differentiation (+Dox) for WT H3 and WT H3K4M cells treated with +DMSO (control) or +DNMT1 inhibitor (DNMT1Ì). Cell numbers are represented in the Iog10 scale, n = 3. E. Interaction plot representing the day 4 over day 0 ratio (Iog2) of cell numbers from the WT H3 or WT H3K4M EpiLC four-day growth assay (+Dox) treated with +DMSO (control) or +DNMT1 inhibitor (DNMT1Ì). * The grey line represents expected results if effects from 5meC and H3K4me depletion were additive, n = 3. F. Four-day growth assay during EpiLC differentiation (+Dox) for RbBP5-FKBP lines treated with +DMSO (control) or +dTAG-13. Cell numbers are represented in the log 10 scale, n = 3. G. Fluorescent reporter (Ruby) signal assessed by flow cytometry in H3 and H3K4M cell lines grown in 2i+VitC (day 0) or in EpiLCs (day 2) (+Dox). Ruby signal is represented in the log 10 scale. A random selection of 1000 individual cells for each cell line is shown. The median is represented. H. Immunoblot showing transgene (HA or H3K4M) expression in cells grown in 2i+VitC (day 0; -/+Dox) or EpiLCs (day 2; +Dox). I. Four-day growth assay during EpiLC differentiation (+Dox). Cell lines are grown either in the presence of a Mock or of a PRC2 inhibitor (PRC2i). Cell numbers are represented in the log 10 scale, n = 3. J. Interaction plot representing the day 4 over day 0 ratio (Iog2) of cell numbers from EpiLC four-day growth assay (+Dox) treated with ÷Mock or+PRC2i. * The grey line represents expected results if effects from PRC2 inhibition compared to Mock conditions in TKO H3 cells were additive in TKO H3K4M cells, n = 3. **A-F, I, J:** Data are shown as mean ± s.e.m. of replicates from independent experiments. P-values were calculated using an unpaired Student’s t-test on day 4 cell counts **(D, F, I),** or a two-way ANOVA test **(E). \*** P≤0.05, ** P≤0.01, *** P≤0.001, **** P≤0.0001.

**Supplemental Figure 6.**
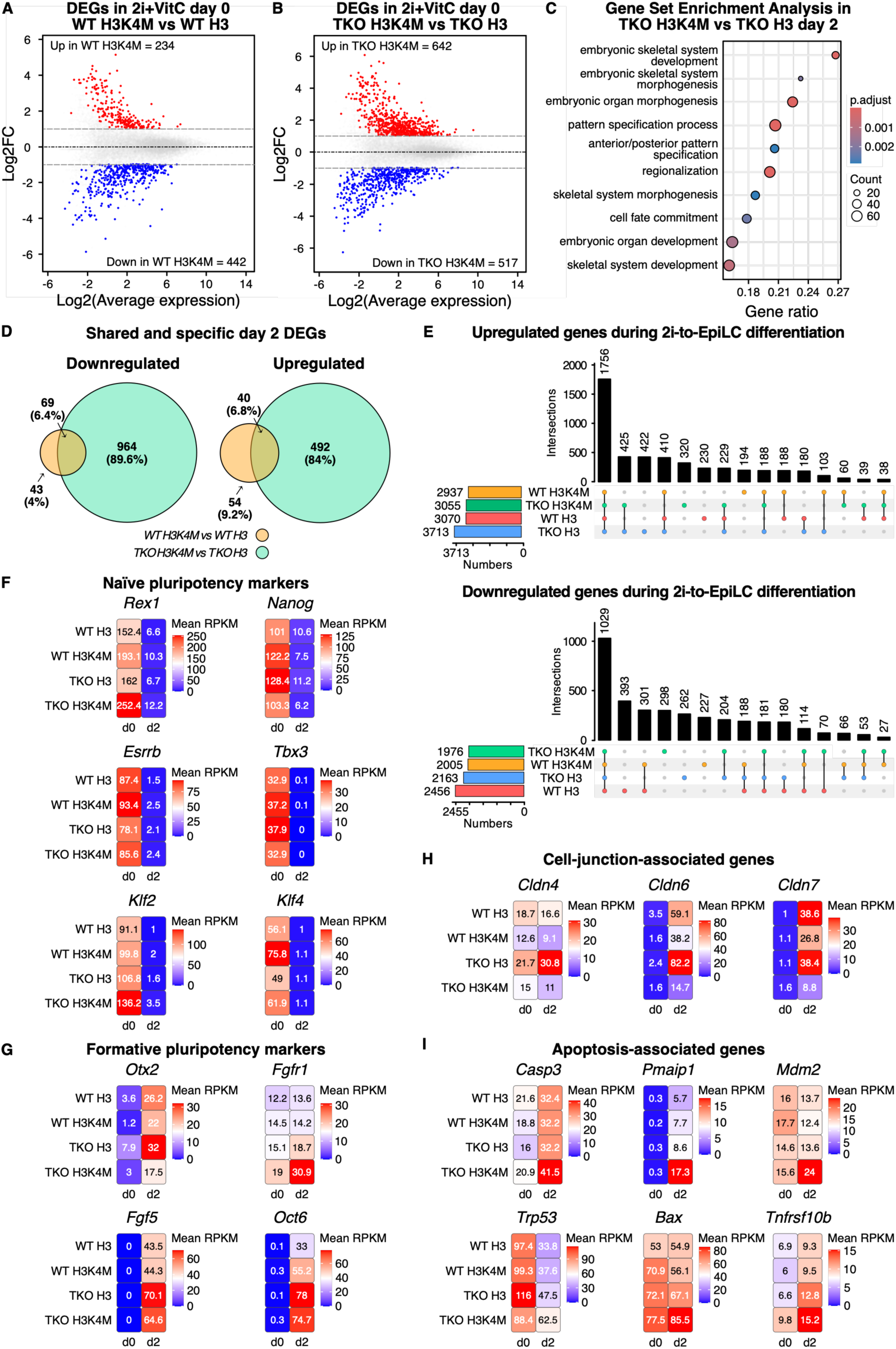
H3K4M-associated expression defects are aggravated in TKO H3K4M cells. **A, B.** MA plot of differential gene expression (|log2FC| > 1 and adjusted p-value (t-test) < 0.05) in ESCs grown in 2i+VitC (day 0) for WT H3K4M vs WT H3 (A) and TKO H3K4M vs TKO H3 (B) comparisons. The average expression is represented as Iog2. Numbers of significantly up- and downregulated genes are indicated. **C.** Gene Set Enrichment Analysis (GSEA) in day 2 EpiLCs for TKO H3K4M vs TKO H3 comparison. D. Venn diagram representing the shared and specific number of down-(left) and upregulated (right) DEGs between WT H3K4M vs WT H3 and TKO H3K4M vs TKO H3 comparisons in day 2 embryoid bodies. E. UpSet plots representing the numbers of shared and unique upregulated (top) and downregulated (bottom) DEGs in EpiLCs (day 2) vs 2i+VitC-grown ESCs (day 0) comparison for each cell line. F-l. Heatmap showing mean expression levels (RPKM) of naive pluripotency markers (F), formative pluripotency markers (G), cell-junction-associated genes (H) and apoptosis-associated genes (I) in 2i+VitC-grown ESCs (day 0) and in EpiLCs (day 2). The mean of two replicates is shown for each cell line and each time point.

**Supplemental Figure 7.**
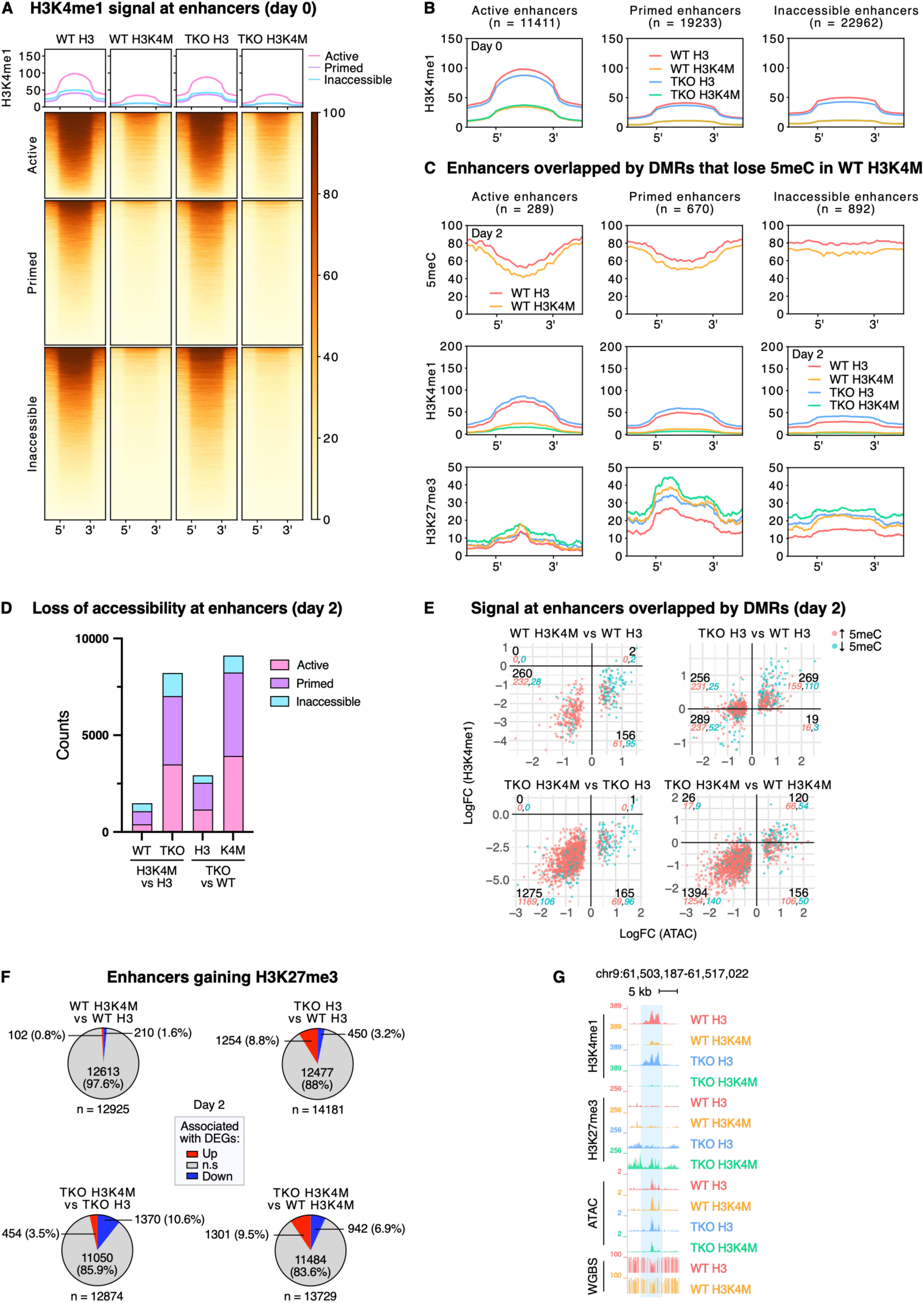
Chromatin and transcriptional consequences of H3K4me1 loss at enhancers in naive ESCs and formative EpiLCs. **A, B.** Heatmap **(A)** and profile plots **(B)** of normalized H3K4me1 signal at enhancers and 1 kb upstream and downstream in naive ESCs (day 0). Enhancers are clustered by category. The mean of two replicates is shown. **C.** Profile plots of 5meC, normalized H3K4me1 and normalized H3K27me3 signals at enhancer DMRs losing 5meC in WT H3K4M vs WT H3, and 1 kb upstream and downstream. Enhancers are clustered by category and signal is shown for all cell lines in EpiLCs (day 2). The mean of two replicates is shown. D. Barplots representing the number of enhancers, clustered by category, losing accessibility in the first cell line of each comparison (FDR < 0.05), in EpiLCs (day 2). **E.** Scatterplots for all comparisons showing the fold-change of ATAC signal (Iog2) vs the fold-change of H3K4me1 signal (Iog2) at enhancers overlapped by DMRs that gain (red) or lose (blue) 5meC in the WT H3K4M vs WT H3 comparison. Only enhancers with statistically-significant differential accessibility (FDR < 0.05) are depicted. Negative values indicate signal loss in the first cell line of each comparison, whereas positive values indicate signal gain in the first cell line. The number of enhancers (black), as well as the number of enhancer DMRs gaining 5meC (red) or losing 5meC (blue) are indicated in each quadrant. F. Pie charts representing the number of enhancers gaining H3K27me3 in the first cell line of each comparison in EpiLCs (day 2), and the number and percentages of this enhancer set associated with significantly up-(red) or downregulated (blue) genes. **G.** UCSC Genome Browser screenshot of a putative active enhancer (highlighted region) in EpiLCs (day 2). From top to bottom, normalized H3K4me1 and H3K27me3 CUT&Tag signals as well as ATAC-seq signals are shown for all cell lines, and 5meC levels in WT lines.

**Supplemental Figure 8.**
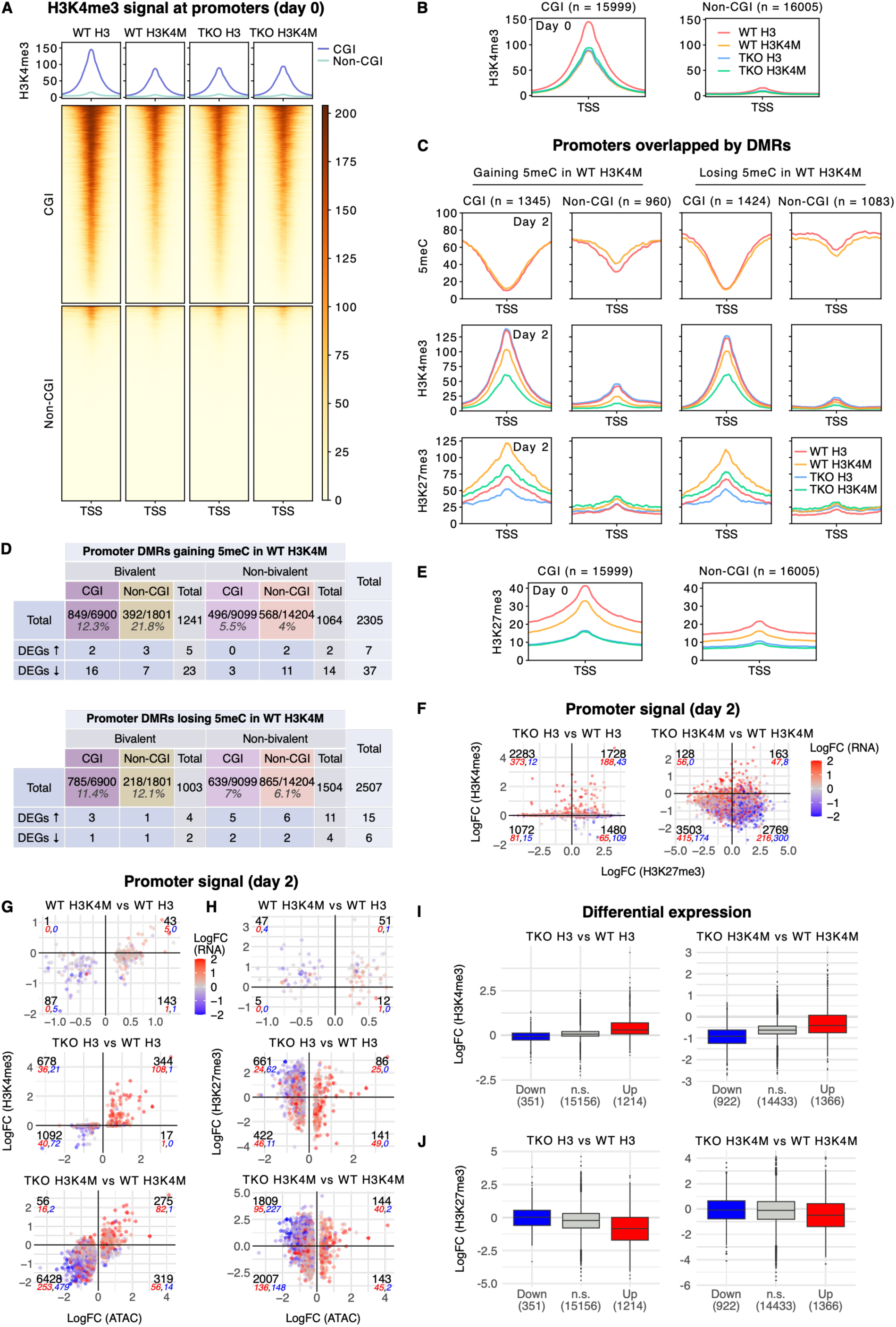
H3K4M expression leads to promoter-associated chromatin changes following H3K4me3 depletion. **A, B.** Heatmap **(A)** and profile plots **(B)** of normalized H3K4me3 signal at CGI and non-CGI promoters (± 2 kb around the TSS) in naive ESCs (day 0). The mean of two replicates is shown. **C.** Profile plots of 5meC, normalized H3K4me3 and normalized H3K27me3 signals at promoters (± 2 kb around the TSS) overlapped by Differentially Methylated Regions (DMRs) that gain (left) or lose (right) 5meC in WT H3K4M vs WT H3. Promoters are clustered by CGI or non-CGI and signal is shown in EpiLCs (day 2). The mean of two replicates is shown. **D.** Table showing the total numbers of DMRs identified at CGI and non-CGI promoters that gain 5meC (top) or lose 5meC (bottom) in WT H3K4M vs WT H3, as well as the number of DMRs associated with up- or downregulated DEGs in WT H3K4M vs WT H3 comparison. **E.** Profile plots of normalized H3K27me3 signal at CGI and non-CGI promoters (± 2 kb around the TSS) in naive ESCs (day 0). The mean of two replicates is shown. F. Scatterplots showing the fold-change of H3K27me3 signal (Iog2) vs the fold-change of H3K4me3 signal (Iog2) at promoters in the different comparisons, colored by the fold-change of associated gene expression (log2(RPKM)) in EpiLCs (day 2). **G, H.** Scatterplots showing the fold-change of ATAC signal (Iog2) vs the fold-change of H3K4me3 **(G)** or H3K27me3 (H) signal (Iog2) at promoters for all comparisons except TKO H3K4M vs TKO H3, colored by the fold-change of associated gene expression (log2(RPKM)) in EpiLCs (day 2). Only promoters with statistically-significant differential accessibility (FDR < 0.05) are depicted. For F-H, negative values indicate signal loss in the first cell line of each comparison, whereas positive values indicate signal gain in the first cell line. The number of promoters (black) as well as the number of associated significantly upregulated (red) or downregulated (blue) DEGs are indicated in each quadrant. I, J. Boxplots showing fold-change of H3K4me3 signal (Iog2) (I) and H3K27me3 signal (J) at promoters of downregulated (Down), non-significant (n.s.) or upregulated (Up) genes (|log2FC| > 1 and adjusted p-value (t-test) < 0.05) in TKO lines in the TKO vs WT comparisons in EpiLCs (day 2).

**Supplemental Figure 9.**
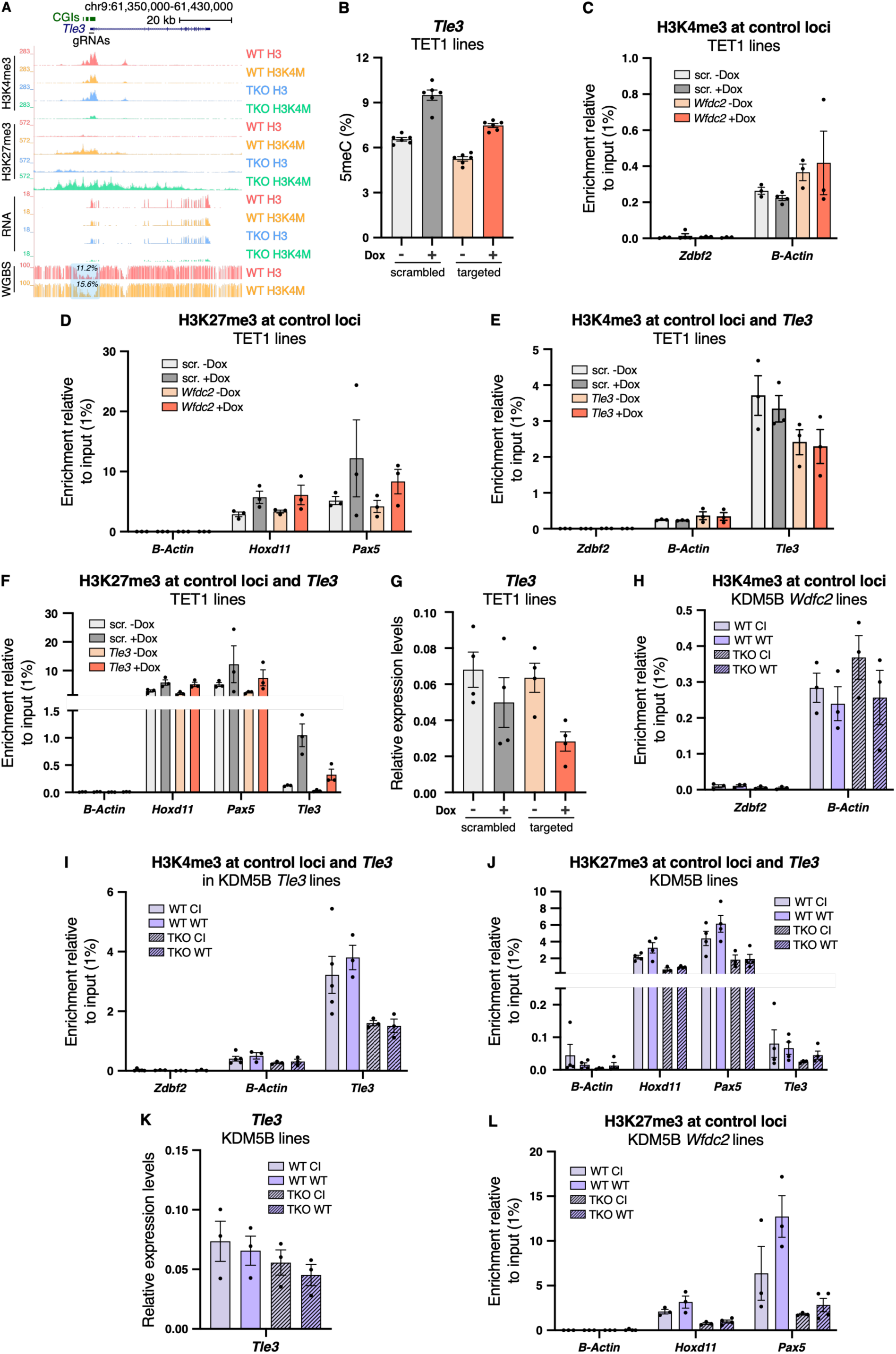
Targeted loss of 5meC or H3K4me3 result in local chromatin changes at *Wfdc2* and *Tle3* promoters. **A.** UCSC Genome Browser screenshot of the *Tle3* locus in all cell lines in EpiLCs (day 2). CGIs and the region targeted by gRNAs are indicated. From top to bottom, normalized H3K4me3 and H3K27me3 CUT&Tag signals, RNA-seq signal, and 5meC levels in WT lines are shown. Average 5meC levels over the highlighted region are indicated in WT lines. B. 5meC levels assessed by bisulfite-pyrosequencing at *Tle3* promoter in TET1 epigenome editing lines. Assay was performed in EpiLCs (day 2) with uninduced or induced H3K4M expression (-/+ Dox). Each data point represents the average 5meC level over the pyrosequencing amplicon (composed of six CpGs) for each replicate. Replicates and their mean are shown, n = 6. **C, D.** H3K4me3 **(C)** and H3K27me3 **(D)** levels assessed by ChlP-qPCR at control loci in TET1 epigenome editing lines targeted at *Wfdc2* in EpiLCs (day 2) with uninduced or induced H3K4M expression (-/+ Dox). For H3K4me3 **(C),** *Zdbf2* represents a negative control locus, and *β-Actin* a positive control locus. For H3K27me3 **(D),** *β-Actin* represents a negative control locus, while *Hoxd11* and *Pax5* represent positive control loci. Enrichment is calculated relative to input (1%). Replicates and their mean are shown, n = 3 or 4. E, F. H3K4me3 (E) and H3K27me3 (F) levels assessed by ChlP-qPCR at the *Tle3* promoter (for (F): left) and control loci (for (F): right) in TET1 epigenome editing lines in EpiLCs (day 2) with uninduced or induced H3K4M expression (-/+ Dox). Enrichment is calculated relative to input (1%). Replicates and their mean are shown, n = 3. **G.** *Tle3* expression assessed by RT-qPCR in TET1 epigenome editing lines in EpiLCs (day 2) with uninduced or induced H3K4M expression (-/+ Dox). Relative expression levels are normalized to the average Ct of the two housekeeping genes *Rrm2* and *RpIpO* using the ΔCt method, n = 4. H. H3K4me3 levels assessed by ChlP-qPCR at control loci in KDM5B *Wfdc2* epigenome editing lines in EpiLCs (day 2). Enrichment is calculated relative to input (1%). Replicates and their mean are shown, n = 3. I. H3K4me3 levels assessed by ChlP-qPCR at the *Tle3* promoter and control loci in KDM5B epigenome editing lines in EpiLCs (day 2). Enrichment is calculated relative to input (1%). Replicates and their mean are shown, n = 3 or 5. J. H3K27me3 levels assessed by ChlP-qPCR at *Tle3* promoter and control loci in KDM5B epigenome editing lines in EpiLCs (day 2). Enrichment is calculated relative to input (1%). Replicates and their mean are shown, n = 4, except TKO KDM5B Cl, n = 3. **K.** *Tle3* expression assessed by RT-qPCR in KDM5B epigenome editing lines in EpiLCs (day 2). Relative expression levels are normalized to the average Ct of the two housekeeping genes *Rrm2* and *RpIpO* using the ΔCt method, n = 3. **L.** H3K27me3 levels assessed by ChlP-qPCR at control loci in KDM5B epigenome editing lines targeting *Wfdc2* in EpiLCs (day 2). Enrichment is calculated relative to input (1%). Replicates and their mean are shown, n = 3 or 4. B-L: Data are shown as mean ± s.e.m. of replicates from independent experiments.

## Supplemental Tables

*Supplemental Table 1*: Guide RNAs and oligo sequences

*Supplemental Table 2*: List of antibodies

